# Benchmarking splice variant prediction algorithms using massively parallel splicing assays

**DOI:** 10.1101/2023.05.04.539398

**Authors:** Cathy Smith, Jacob O. Kitzman

## Abstract

**Background:** Variants that disrupt mRNA splicing account for a sizable fraction of the pathogenic burden in many genetic disorders, but identifying splice-disruptive variants (SDVs) beyond the essential splice site dinucleotides remains difficult. Computational predictors are often discordant, compounding the challenge of variant interpretation. Because they are primarily validated using clinical variant sets heavily biased to known canonical splice site mutations, it remains unclear how well their performance generalizes.

**Results:** We benchmarked eight widely used splicing effect prediction algorithms, leveraging massively parallel splicing assays (MPSAs) as a source of experimentally determined ground-truth. MPSAs simultaneously assay many variants to nominate candidate SDVs. We compared experimentally measured splicing outcomes with bioinformatic predictions for 3,616 variants in five genes. Algorithms’ concordance with MPSA measurements, and with each other, was lower for exonic than intronic variants, underscoring the difficulty of identifying missense or synonymous SDVs. Deep learning-based predictors trained on gene model annotations achieved the best overall performance at distinguishing disruptive and neutral variants. Controlling for overall call rate genome-wide, SpliceAI and Pangolin also showed superior overall sensitivity for identifying SDVs. Finally, our results highlight two practical considerations when scoring variants genome-wide: finding an optimal score cutoff, and the substantial variability introduced by differences in gene model annotation, and we suggest strategies for optimal splice effect prediction in the face of these issues.

**Conclusion:** SpliceAI and Pangolin showed the best overall performance among predictors tested, however, improvements in splice effect prediction are still needed especially within exons.

## Background

Splicing is the process by which introns are removed during mRNA maturation using sequence information encoded in the primary transcript. Sequence variants which disrupt splicing contribute to the allelic spectrum of many human genetic disorders, and it is estimated that overall, as many as 1 in 3 disease-associated single-nucleotide variants are splice-disruptive^1-7^. Splice-disruptive variants (SDVs) are most readily recognized at the essential splice site dinucleotides (GU/AG for U2-type introns) with many examples across Mendelian disorders^8-12^. SDVs can also occur at several so-called flanking noncanonical positions^13^, which by some estimates outnumber essential splice mutations by several-fold^5,14^.

Variants beyond the splice-site motifs may be similarly disruptive but are more challenging to recognize^15^. For instance, some SDVs disrupt splicing enhancers or silencers, short motifs bound by splicing factors to stimulate or suppress nearby splice sites, to confer additional specificity, and to provide for regulated alternative splicing^16^. These elements are widespread^17^ and maintained by purifying selection^18^, but their grammar is often unclear as they feature partial redundancy and can tolerate some mutations. Nevertheless, variants which disrupt splicing regulatory elements have been implicated in a number of disorders. In one prominent example, spinal muscular atrophy, loss of *SMN1* cannot be fully complemented by its nearly identical paralog *SMN2* due to the loss of an exonic splice enhancer (ESE) in exon 7 of the latter gene^19,20^, a defect which can be therapeutically targeted by antisense oligonucleotides^21^. Synonymous variants, which as a class may be overlooked, may also disrupt existing splice regulatory elements or introduce new ones, as in the case of *ATP6PA2-*associated X-linked parkinsonism^22^.

RNA analysis from patient specimens can provide strong evidence for splice-disruptive variants, and its inclusion in clinical genetic testing can improve diagnostic yield^5,23-26^. However, advance knowledge of the affected gene is necessary for targeted RT-PCR analysis, while RNA-seq-based tests are not yet widespread^27,28^, and both rely upon sufficient expression in the blood or other clinically accessible tissues for detection. Therefore, a need remains for reliable *in silico* prediction of SDVs during genetic testing, and a diverse array of algorithms have been developed to this end. For instance, S-Cap^29^ and SQUIRLS^30^ implement classifiers that use features such as motif models of splice sites, kmer scores for splice regulatory elements, and evolutionary sequence conservation, and are trained on sets of benign and pathogenic clinical variants. Numerous recent algorithms use deep learning to predict splice site likelihoods directly from the primary sequence; SDVs can then be detected by comparing predictions for wild-type and mutant sequence. Rather than training with clinical variant sets, SpliceAI^31^ and Pangolin^32^ are trained using gene model annotations to label each genomic position as true/false based on whether it appears as an acceptor or donor in a known transcript. SPANR^33^ uses the primary sequence to predict percent spliced in (PSI) measurements with training data provided by RNA-seq measurements. HAL^34^ takes a distinct approach by training on a library of randomized sequences and their experimentally observed splicing patterns, while MMSplice^35^ combines the training data from HAL with features derived from primary sequence and additional modules trained on annotated splice sites and clinical variants. Finally, ConSpliceML^36^ is a metaclassifier that combines SQUIRLS and SpliceAI scores with a population-based constraint metric measuring the regional depletion of predicted splice-disruptive variants among apparently healthy adults in population databases.

Given the proliferation of splicing predictors and their utility in variant interpretation, it is important to understand their performance characteristics. Previous comparisons have suggested that overall, SpliceAI represents the state-of-the art, with several other algorithms including MMSplice, SQUIRLS, and ConSpliceML showing competitive or in some cases better performance^30,32,36-42^. However, benchmarking efforts to date primarily relied upon curated sets of clinical variants^30,36,37,39-42^, which are strongly enriched for canonical splice site mutations^33,37,42-45^ likely due to the relative ease of their classification. This leaves open the question of how well these tools’ performance may generalize, and whether certain tools may excel in particular contexts (e.g., for exonic cryptic splice activating mutations). A further challenge is that some of these tools’ training data may partially overlap with benchmarking validation sets which risks circularity if overlapping variants are not carefully identified and removed.

Massively parallel splicing assays (MPSAs) provide an opportunity to benchmark splicing effect predictors entirely orthogonally to clinical and population variant sets. MPSAs measure thousands of variants’ splicing effects in a pooled fashion: cells are transfected with a library of variants cloned into a minigene construct with deep RNA sequencing as a quantitative readout of variants’ splicing outcomes. MPSAs come in several different flavors: broad MPSA screens assess many exons and measure one or a few variants’ effects at each^6,14,46,47^, while saturation screens focus on individual exons^48-53^ or motifs^34,54^ and measure the effects of every possible point variant within each target. Two broad MPSA datasets, Vex-seq^46^ and MaPSy^6^, were recently used to benchmark splicing effect predictors as part of the Critical Assessment of Genome Interpretation (CAGI) competition^55^, and another, MFASS^14^, has been used to validate a recent meta-predictor^38^. However, a limitation of benchmarking with broad MPSAs is that they may reflect an exon’s overall properties while lacking the finer resolution to assess different variants within it. For instance, an algorithm could perform well by predicting SDVs within exons with weak splice sites, or with evolutionarily conserved sequence, while failing to distinguish between truly disruptive and neutral variants within each.

Here we leverage saturation MPSAs as a complementary, high-resolution source of benchmarking data to evaluate eight recent and widely used splice predictors. Algorithms using deep learning to model splicing impacts using extensive flanking sequence contexts, SpliceAI and Pangolin, consistently showed the highest agreement with measured splicing effects, while other tools performed well on specific exons or variant types. Even for the best performing tools, predictions were less concordant with measured effects for exonic variants versus intronic ones, indicating a key area of improvement for future algorithms.

## Results

### A validation set of variants and splice effects

We aggregated splicing effect measurements for 2,230 variants from four massively parallel splice assay (MPSA) studies, focusing on saturation screens targeting all single nucleotide variants (SNVs) in and around selected exons^48-50,53^ (**Figure 1A**). We also included 1,386 variants in *BRCA1* from a recent saturation genome editing (SGE) study, in which mutations were introduced to the endogenous locus by CRISPR/Cas9-mediated genome editing, with splicing outcomes similarly measured by RNA sequencing^56^. For contrast with these saturation-scale datasets, we also prepared a more conventional, gene-focused benchmarking dataset by manually curating a set of 296 variants in the tumor suppressor gene *MLH1* from clinical variant databases and literature reports. In sum, this benchmarking dataset contained 3,912 SNVs across 33 exons spanning six genes (**Supplementary Table 1**; **Supplementary Figures 1-5**).

**Figure 1.**
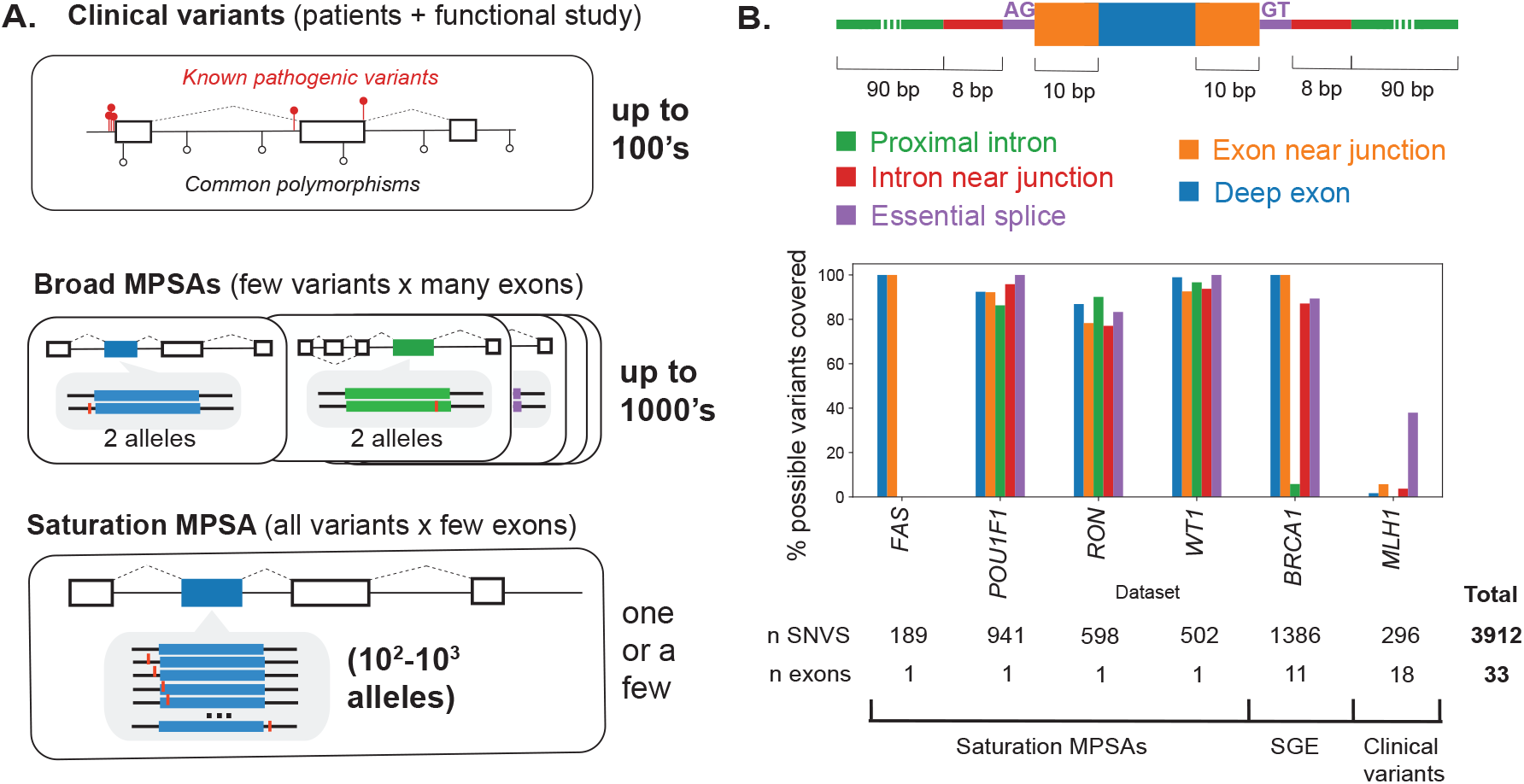
Variants used for splice effect predictor benchmarking. **(A)** Validation sets can be drawn from (top panel) pathogenic clinical variants and, conversely, common polymorphisms in frequently screened disease genes, (middle panel) broadly targeted massively parallel splice assays (MPSAs) interrogating a few variants across many exons, and (bottom panel) saturation MPSAs in which all possible variants are created for a few target exons. **(B)** Variant classes defined by exon/intron region and proximity to splice sites (upper), with the percent coverage of the possible SNVs within each variant class (denoted by color) for each dataset in the benchmark set (for *BRCA1*, missense and stop-gain variants were excluded and not counted in the denominator).

As expected, MPSAs measured most of the possible single-nucleotide variants at each target (93.3% of SNVs) with relatively uniform coverage by exon/intron region (**Figure 1B**). From the *BRCA1* SGE study, we retained only intronic or synonymous variants because missense variants’ effects could be mediated via protein alteration, splicing impacts, or both. Targeted exons varied in their robustness to splicing disruption, from *POU1F1* exon 2 (10.2% SDV), to *RON* exon 11 (68.4% SDV; **Supplementary Figure 6**), reflecting both intrinsic differences between exons as well as different procedures for calling SDVs across MPSA studies. In contrast to the high coverage of the mutational space from MPSA and SGE datasets, reported clinical variants only sparsely covered the mutational space (1.6% of the possible SNVs in *MLH1* exons +/- 100 bp) and were heavily biased towards splice sites (59.5% of reported variants within +/-10 bp of a splice site; **Supplementary Figure 7**). Larger clinical variant sets used to train classifiers showed a similar skew: 94.6% of the SQUIRLS training variants^30^ and 88.9% of the pathogenic S-Cap training set^29^ were within +/- 10 bp of a splice sites. Thus, MPSAs offer high coverage without the variant class biases present among clinical variant sets.

### Comparing bioinformatic predictions with MPSA measured effects

We selected eight recent and widely used predictors to evaluate: HAL^34^, S-Cap^29^, MMSplice^35^, SQUIRLS^30^, SPANR^33^, SpliceAI^31^, Pangolin^32^, and ConSpliceML^36^. Most variants (93.1%) were scored by all tools except HAL and S-Cap (which focus on only exonic variants and synonymous/proximal intronic variants, respectively). Algorithms’ predictions were only modestly correlated with each other (median pairwise Pearson *r* between absolute values of tools’ scores = 0.58, range: 0.04 to 0.97; **Supplementary Figure 8**). One exception was Pangolin and SpliceAI, which share similar model architectures and training sets, and were almost perfectly correlated with each other (*r* = 0.97). These two were also strongly correlated with MMSplice (*r* = 0.81 and 0.80 respectively). The pattern of modest agreement across tools was not specific to exons and variants tested by MPSAs: we observed a similar degree of correlation between tools’ scores across a background set of randomly sampled genomic SNVs (median pairwise *r* = 0.60; range: 0.07 to 0.93; **Supplementary Figure 8** and **Methods**). Concordance between algorithms was notably lower within exons, both in the MPSA benchmarking set variants (median pairwise *r* = 0.43) and random background set variants (median *r* = 0.39).

Agreement between predictors’ scores and experimentally measured effects also varied widely by algorithm and MPSA dataset, but were similarly modest, with a median Pearson’s *r* of 0.53 (range of -0.06 to 0.85; **Supplementary Figure 9**). Even for a single algorithm and exon, substantial regional variability in concordance was evident (**Figure 2**). For instance, at *POU1F1* exon 2, every tool other than S-Cap recapitulated the strong constraint observed by MPSA at the donor region. By contrast, at a putative exonic splicing silencer (ESS) near the alternative beta acceptor, algorithmic and measured effects were much less concordant, reflecting the difficulty of modeling variant effects outside canonical splicing motifs.

**Figure 2.**
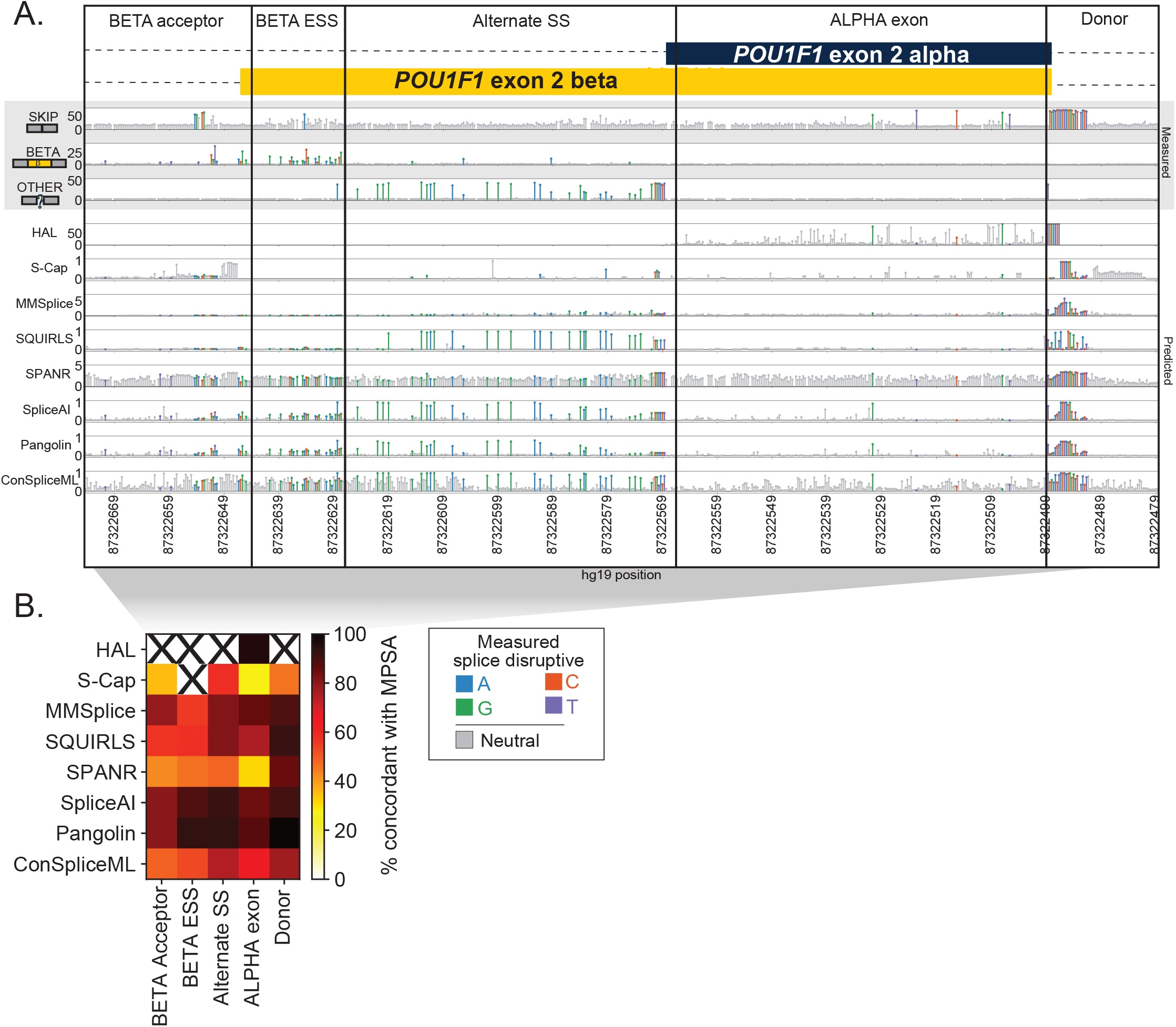
Agreement between predictors and experiments varies by gene region. **(A)** Splicing effect tracks at alternate isoforms beta and alpha from a published MPSA^53^ of *POU1F1* exon 2. Upper panel tracks (gray background) show MPSA-measured percent of exon skipping, beta exon inclusion, and other (non-alpha/beta/skip) isoform usage; lower tracks show scores from bioinformatic predictors by position. Each lollipop denotes one variant, shaded by effect in MPSA (gray: neutral, colors: SDVs, shaded by mutant base). The exon and flanking introns are split by region. **(B)** Heatmap showing concordance between each algorithm’s binary classification of variants as SDV/neutral versus those of the MPSAs, at the optimal score threshold for each algorithm. Concordance is shown per algorithm (row) for each region (column). Regions with <10 scored variants are omitted (‘X’ symbols).

To systematically benchmark each predictor, we treated the splicing status from the experimental assays and curated clinical variant set as ground truth. We quantified the ability of each predictor to distinguish between the splice disruptive (*n*=1,060) and neutral (*n*=2,852) variants in the benchmark set by taking the area under the precision-recall curve (prAUC) per classifier/gene (**Figure 3A**). We next asked if classifiers’ performance differed by variant type and location. Algorithms consistently performed better for intronic than for exonic variants (median prAUC for introns: 0.773; for exons: 0.419; **Figure 3B**), despite a similar proportion of SDVs in exons and introns (28.4% and 25.9% SDV, respectively). This difference persisted even when removing canonical splice dinucleotide variants (**Supplementary Figure 10**). More finely subdividing the benchmark variant set by regions (defined as in **Figure 1B)** demonstrated that performance suffers further from splice sites, where the overall load of SDVs is lower (**Supplementary Figure 11**). To summarize overall performance, we counted the number of instances in which each predictor either had the highest prAUC or was within the 95% confidence interval of the winning tool’s prAUC (**Figure 3C)**. Every tool scored well for at least one dataset or variant class, but Pangolin and SpliceAI had the best performance most frequently (7 and 3 datasets/variant classes, respectively).

**Figure 3.**
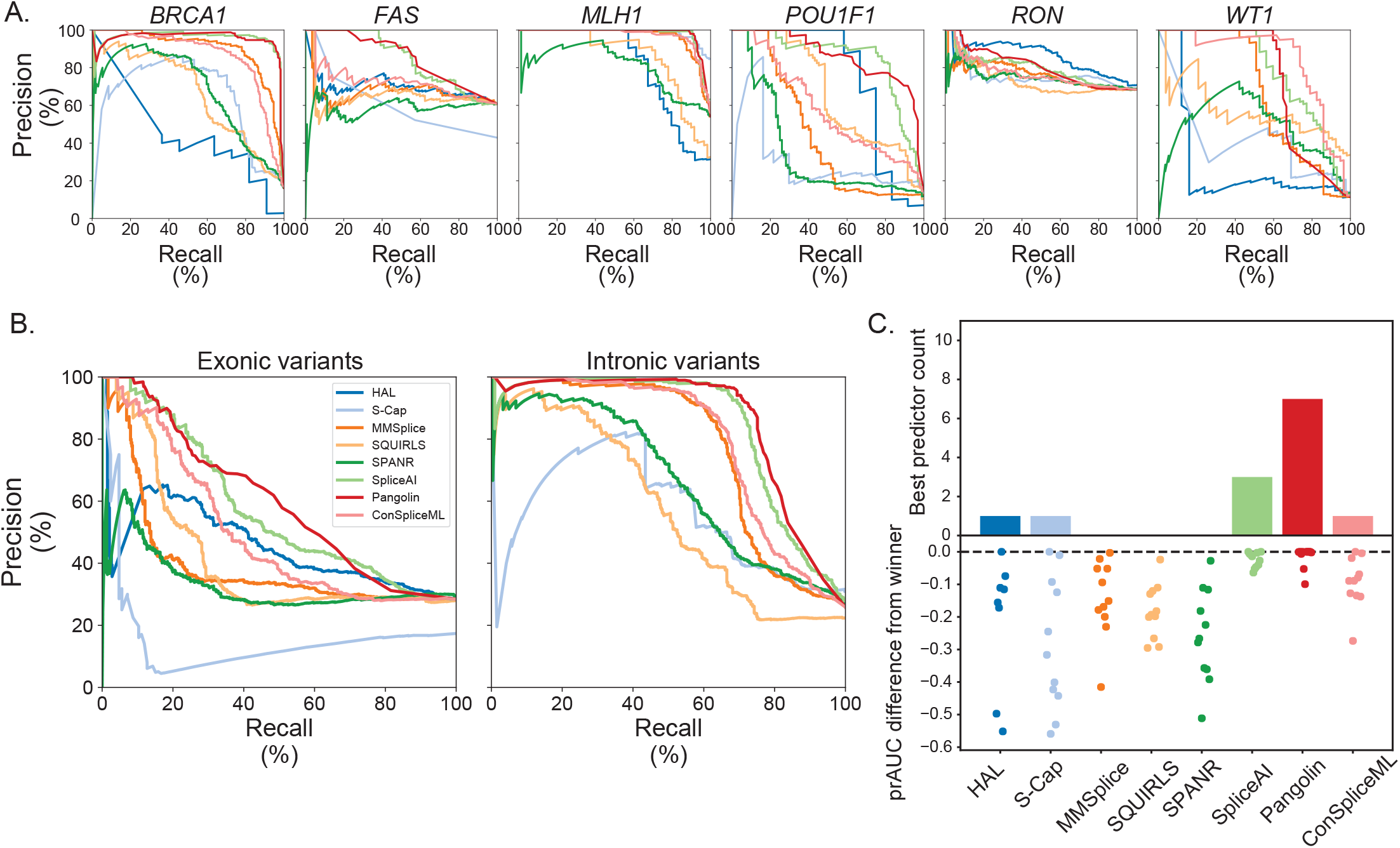
Splice effect predictors’ classification performance on benchmark variants. **(A)** Precision-recall curves showing algorithms’ performance distinguishing SDVs and splicing-neutral variants in each dataset. **(B)** Precision-recall curves of tools’ performance differentiating SDVs and splice neutral variants in exons (left) and introns (right). **(C)** Top panel: tally, for each algorithm, of the number of individual datasets and variant classes (defined as in **Figure 1B**) for which that algorithm had the highest prAUC or was within the 95% confidence interval of the winning tool. Bottom panel: signed difference between the winning tool’s prAUC and a given tool’s prAUC; each dot corresponds to an individual dataset or variant class.

### Benchmarking in the context of genome-wide prediction

In practice, a splicing effect predictor must sensitively identify SDVs while maintaining a low false positive rate across many thousands of variants identified in an individual genome. We therefore evaluated each tool’s sensitivity for SDVs within our benchmark set, as a function of its genome-wide SDV call rate. We used a background set of 500,000 simulated SNVs drawn at random from in or near (+/- 100 bp) internal protein-coding exons **(Supplementary Figure 12; Supplementary Table 1)**. We scored these background SNVs with each tool and computed the fraction of the background set called as SDV as a function of the tool-specific score threshold. Although the true splice-disruptive fraction of these background variants is unknown, we normalized algorithms to each other by taking, for each algorithm, the score threshold at which it called an equal fraction (e.g., 10%) of the genomic background set as SDV **(Supplementary Table 2)**. We then computed the sensitivity across the benchmark-set SDVs using this score threshold and termed this the ‘transcriptome-normalized sensitivity’. Taking SpliceAI as an example, at a threshold of deltaMax≥0.06, 10% of the background set is called as SDV. Applying the same threshold (deltaMax≥0.06) to *BRCA1* SGE variants in the benchmark set, SpliceAI reaches 98.2% sensitivity and 80.7% specificity (**Figure 4A**).

**Figure 4.**
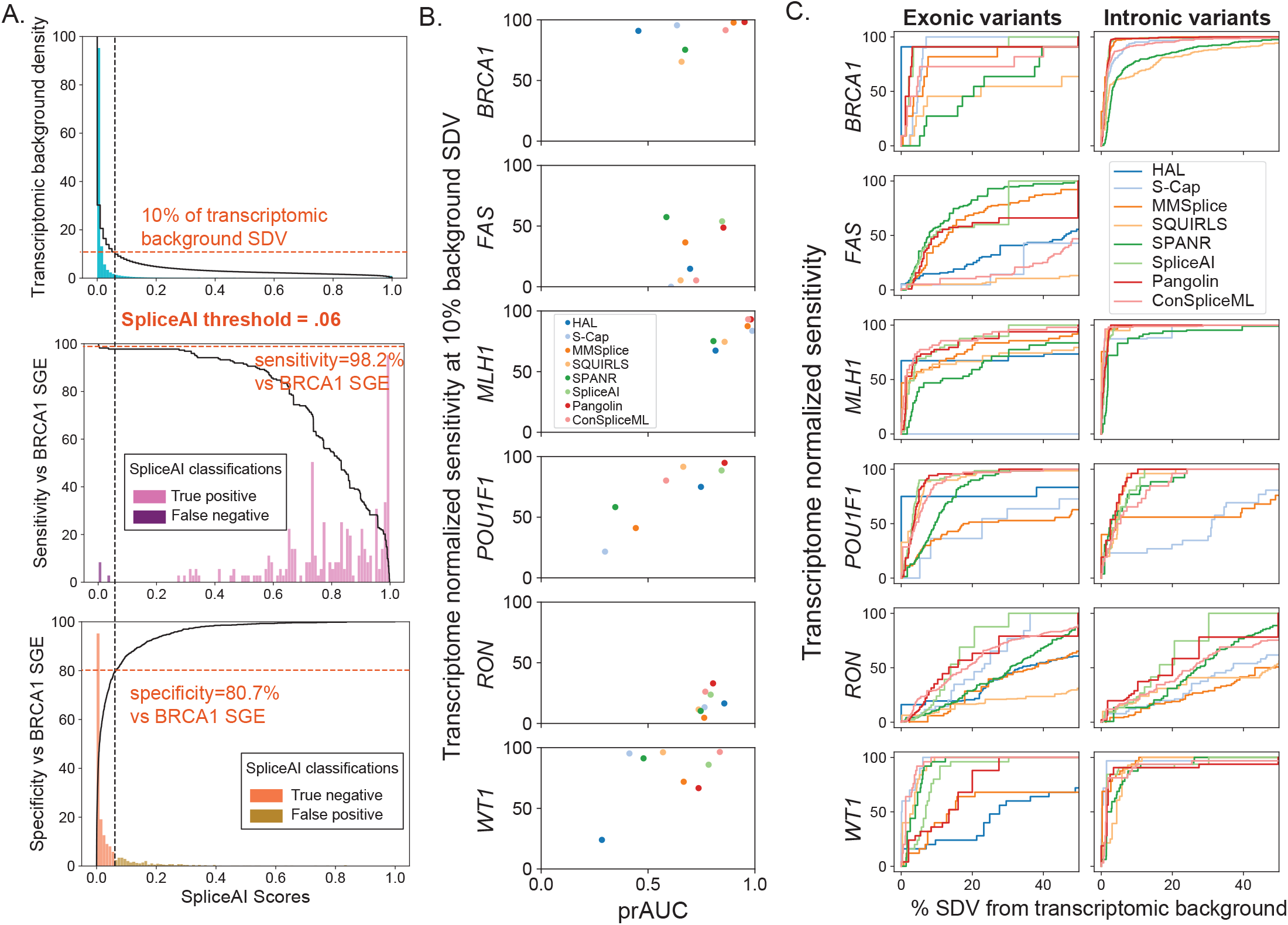
Transcriptome-normalized sensitivity. **(A)** Example shown for SpliceAI. Upper panel shows SpliceAI scores for the 500,000 background set variants (teal histogram) and the cumulative fraction (black line) of variants above a given score threshold (10% of background set variants with deltaMax≥0.06). Below, histograms of SpliceAI scores for *BRCA1* SGE benchmark variants, either SDVs (middle) or splicing-neutral variants (bottom) and the resulting transcriptome-normalized sensitivity and specificity at a deltaMax cutoff of 0.06. **(B**) Transcriptome-normalized sensitivity (at 10% background set SDV) versus within-benchmark variant set prAUC by benchmarked dataset. **(C)** Transcriptome-normalized sensitivity on benchmark exonic and intronic variants plotted as a function of the percent of the background variant set called SDV.

We repeated this process, using for each algorithm the score threshold at which 10% of the background set was called as SDV, and applying this threshold to the benchmark set **(Supplementary Table 2)**. Transcriptome-normalized sensitivity varied widely between algorithms, but SpliceAI, ConSpliceML, and Pangolin emerged as consistent leaders (median across datasets of 87.3%, 85.8%, 79.9%, respectively). Mirroring the results seen entirely within the benchmarking variant set (**Figure 3**), median transcriptome-normalized sensitivity was lower for exonic vs intronic variants for all tools examined by an average of 36.9%, and the same pattern remained after removing intronic variants at essential splice sites. These results were not specific to the transcriptome-wide threshold of 10%: the same three algorithms scored highly for thresholds at which 5% or 20% of the background set scored as SDV **(Supplementary Table 2)**. Performance also varied by exon target (**Figure 4B**); for example, many of the SDVs in *FAS* exon 6 and *RON* exon 11 were not detected by any algorithm at a threshold which would classify 10% of the background set as SDV. The effects measured by MPSAs in these specific exons may be particularly subtle, posing difficult targets for prediction, and suggesting that existing tools may need scoring thresholds tuned to specific exons or variant regions. Finally, we quantified the transcriptome-normalized sensitivity as a function of percent of the background set called SDV, and calculated the area under the resulting curve (analogous to the prAUC statistic), showing the tradeoff between benchmark SDV recall and genome-wide SDV rate, and again highlighting consistently lower performance within exons across algorithms and datasets (**Figure 4C; Supplementary Table 2**).

### Determining optimal score cutoffs

Integrating splice effect predictors into variant interpretation pipelines requires a pre-determined score threshold beyond which variants are deemed disruptive. We explored whether our benchmarking efforts could inform this by identifying the score threshold that maximized the Youden’s J statistic (*J*=sensitivity+specificity-1; **Supplementary Table 3)**. For each algorithm, we first identified optimal score thresholds on each dataset individually to explore differences across genes and exons. For most tools we evaluated, ideal thresholds varied considerably across exons, regions, and variant classes, such that a threshold derived from one was suboptimal for others (**Figure 5**). For some tools, including HAL and ConSpliceML, thresholds optimized on individual datasets spanned nearly the tools’ entire range of scores, while for others such as SQUIRLS, SpliceAI, and Pangolin, the optimal thresholds were less variable. For the tools with consistently high classification performance and transcriptome-normalized sensitivity – SpliceAI, Pangolin, and ConSpliceML (**Figures 3 and 4**) – we found that the optimal thresholds were usually lower than the threshold recommended by the tools’ authors, largely consistent with conclusions of other previous benchmarking efforts^39,40,42,43^. Optimal thresholds also differed by variant class, suggesting that tuning cutoffs by variants’ annotated effects, like those implemented in S-Cap, may offer some improvement for classification accuracy on variants genome-wide.

**Figure 5.**
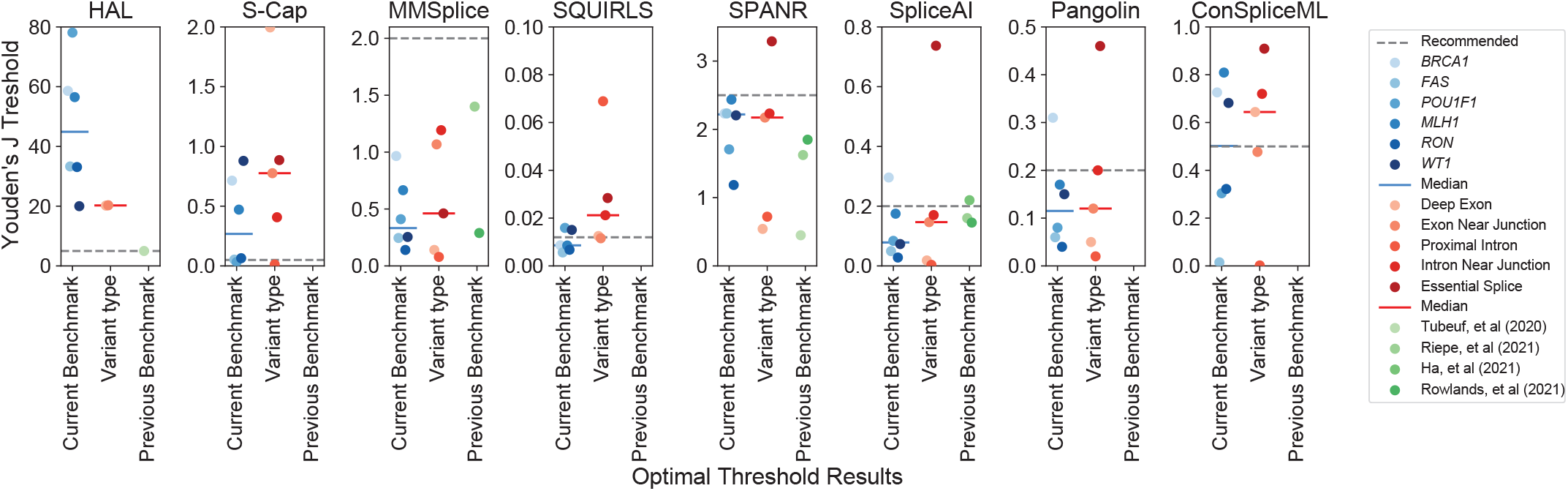
Optimal thresholds to classify splice disruptive variants. Optimal score thresholds (y-axis) for each algorithm across each benchmark variant dataset (blue points), by variant type (red points), and compared to previous reports (green points). Dashed gray line shows tool developers’ recommended thresholds. Solid lines indicate medians.

### Variant effects at alternative splice sites

Alternative splicing can present challenges for variant effect prediction. Many tools require gene model annotation as an input, including four of the eight tested here (MMSplice, SQUIRLS, SpliceAI, and Pangolin). Effect predictions for individual variants may be influenced by the inclusion or exclusion of nearby alternative isoforms in these annotations. For instance, SpliceAI and Pangolin by default apply a mask which suppresses scores from variants that either strengthen known splice sites or weaken unannotated splice sites under the assumption that neither would be deleterious. Although masking may be a useful step to reduce the number of high-scoring variants genome-wide, it requires the provided annotation to be complete and assumes there is no functional sensitivity to the relative balance among alternative splice forms.

We examined the effects of annotation choices and masking options at the two alternatively spliced exons in our benchmark variant set. In the first, *POU1F1*, two functionally distinct isoforms (alpha and beta) result from a pair of competing acceptors at exon 2. Alpha encodes a robust transactivator and normally accounts for ≥97% of *POU1F1* expression in the human pituitary^57-60^. Beta exhibits dominant negative activity, and SDVs that increase its expression cause combined pituitary hormone deficiency^53,61^. We focused on SpliceAI, in which the default annotation file includes only the alpha transcript. Predictions were broadly similar after updating annotations to include only the beta isoform or to include both: 13.8% (*n*=130/941) and 10.5% (*n*=99/941) of the variants, respectively, changed classifications compared to SpliceAI run with default annotations (each at an SDV cutoff of deltaMax≥0.08 which was optimal across the dataset; **Supplementary Figure 13**). Among these were several pathogenic SDVs including c.143-5A>G which is associated with combined pituitary hormone deficiency (CPHD)^62^, scored as highly disruptive by MPSA^53^, and was validated *in vivo* by a mouse model^63^. With the default annotations (alpha isoform only) and when including both isoforms, SpliceAI scores c.143-5A>G as disruptive (deltaMax=0.21 and 0.16, respectively). However, when only the beta isoform is included, this variant is predicted neutral (deltaMax=0). A similar pattern emerged at a cluster of six pathogenic SDVs which disrupt a putative exonic splicing silencer which normally suppresses beta isoform expression^53^. Therefore, counterintuitively, pathogenic SDVs which act by increasing beta isoform usage go undetected when using annotation specific to that isoform.

The choice of canonical transcript may be less clear when alternative isoforms’ expression are more evenly balanced, as in the case of *WT1*, a key kidney and urogenital transcription factor gene^64^ covered by our benchmarking set. Exon 9 of *WT1* has two isoforms, KTS+ and KTS-, named for the additional three amino acids included when the downstream donor is used^65,66^. In the healthy kidney, KTS+ and KTS- are expressed at a 2:1 ratio^67,68^. Decreases in this ratio cause the rare glomerulopathy Frasier’s Syndrome^67-69^, while increases are associated with differences in sexual development (DSD)^70^. We ran SpliceAI using annotations including KTS+ alone (its default), KTS-alone, and with both isoforms (**Supplementary Figure 14**). A cluster of variants, including one associated with DSD near the unannotated KTS-donor^70^ (c.1437A>G), appear to weaken that donor but are masked because the KTS-donor is absent from the default annotations. Conversely, another variant (c.1447+3G>A) associated with DSD appears to increase the KTS+/KTS-ratio but is also masked because it strengthens the annotated KTS+ donor (deltaMax=0 with default annotation), and similarly scores as neutral when the annotation is updated to include both isoforms (deltaMax=0.02). That variant scores somewhat more highly (deltaMax=0.12) when only the KTS-annotation is used, but that in turn results in failure to capture several known Frasier’s Syndrome pathogenic variants near the KTS+ donor^48,67,68,71-74^. This case illustrates that predictors can fail even when all functionally relevant isoforms are included, because masking may suppress SDVs for which the pathogenic effects result from strengthening of an annotated splice sites and disrupting the balance between alternative isoforms. This challenge was not specific to SpliceAI; for instance, Pangolin also showed poor recovery of KTS-SDVs (only 25% correctly predicted) due to a similar masking operation.

*POU1F1* and *WT1* do not represent exceptional cases. Among RNA-seq junction usage data from the GTEx Consortium^60^, we estimate 18.0% of all protein coding genes (*n*=3,571/19,817 genes) have at least one alternate splice site that is expressed and at least modestly used (≥20% PSI) in at least one tissue, yet is absent from SpliceAI default annotations (**Supplementary Figure 15**). One of these is *FGFR2*, a tyrosine kinase gene with key roles in craniofacial development^75,76,77^. Mutually exclusive inclusion of its exons IIIb and IIIc results in two isoforms with different ligand specificites^75,76,78^, and disruption of exon IIIc splicing causes Crouzon, Apert, and Pfeiffer Syndromes, which share overlapping features including craniosynostosis (premature cranial suture fusion)^79-82^. Pathogenic variants cluster near exon IIIc splice sites and at a synonymous site that activates cryptic donor usage within the exon^79,81-92^ (**Supplementary Figure 16**). The default annotation excludes exon IIIc, causing all four pathogenic variants at its acceptor to be scored splice neutral, but when IIIc is included in the annotation, all four are predicted with high confidence (all ≥0.99; **Supplementary Table 1**). Disabling masking in order to capture cases such as these is not a viable option as it greatly reduces overall performance, and drastically increases the number of high-scoring variants which must be reviewed^38^.

## Discussion

We evaluated the performance of eight splice effect predictors using a benchmark set of variants from saturation-level massively parallel splicing assays (MPSAs) across fifteen exons. By holding the sequence context constant for hundreds of variants per exon, these MPSAs afforded an opportunity to systematically evaluate how well each tool could distinguish individual variants’ effects without confounding effects of differences in exons’ overall characteristics. Compared to traditional validation sources such as clinical variant databases which are enriched for essential splice site mutations, these MPSA datasets had more uniform representation of variant types including those for which classification is currently challenging.

Across most exons tested, the deep learning-based tools Pangolin and SpliceAI had the best overall performance. These two were not uniformly superior, however, and other tools excelled on certain datasets. ConSpliceML was comparably sensitive at identifying SDVs within the benchmarking set, while normalizing for genome-wide call rate, and MMSplice performed well for intronic SDVs. Even for the best performing tools, SDVs were more difficult to identify within exons compared to introns, highlighting an area for future improvement. These results are consistent with other recent splice predictor benchmarks using broad MPSAs and clinical variants, which also noted low concordance among tools^38,40^, particularly for exonic variants^42^, and poorer classification performance in exons and with greater distance from splice sites^14,30,31,43,93^. As we found, SpliceAI was often but not always the top performer in these past comparisons^37-42,93^. Together, these our results suggest opportunities for metaclassifiers to better calibrate existing predictors and to leverage each within its strongest domain^36,38^.

A key issue this benchmarking study highlights is the challenge of selecting a scoring threshold for splicing predictors. This may reflect differences in exons’ and genes’ intrinsic vulnerability to SDVs, as a function of factors such as splice site strength^94^ and wildtype exons’ baseline inclusion rates^95^. For instance, most predictors fared poorly on *FAS* exon 6 and *RON* exon 11, both of which are intermediately included at baseline, and so may be more sensitive to splice disruption^95^. For moderately included exons such as these, more lenient thresholds may be required.

Another consideration is that most predictors do not directly model differences in dosage sensitivity between genes. In some genes, SDVs that reduce the abundance of properly spliced mRNA by ∼50% may be tolerated, whereas in more highly dosage sensitive genes, an equivalently splice-disruptive variant would be highly deleterious. The nature of the aberrant splice form is also important to consider; SDVs that increase expression of protein isoforms with dominant negative effects may be deleterious even at a low level of expression as in the case of *POU1F1* exon 2 beta-promoting SDVs. Similarly, while *DNM1* loss of function can result in developmental and epileptic encephalopathies, specific SDVs yield in-frame insertions which act in a dominant negative fashion and cause a particularly severe presentation^96^. Interpreting the results of bioinformatic splice effect predictions may therefore depend upon expert knowledge of the individual genes’ dosage sensitivity which potentially limits the utility of readily computed genome-wide scores. Methods such as ConSpliceML offer a means of inferring such thresholds by modeling the constraint against SDVs among healthy individuals on a regional (e.g., per exon) basis^36^.

Our results also highlight the major influence of gene model annotation, a required input for many splice effect predictors. For two of the MPSA-tested exons in our benchmarking set (*POU1F1* and *WT1*), inclusion of alternate splice forms in the annotation input altered SpliceAI’s predictions across >10% of variants. Using RNA-seq data from GTEx, we conservatively project that this challenge may impact nearly one in every five genes in the human genome. Such annotation changes are inconvenient for end users and are not readily accommodated by some tools. Moreover, they may not be possible when the functionally relevant isoforms are not known in advance. Using the most comprehensive annotation set is not a universal fix, as illustrated by *POU1F1*, where it resulted in poorer concordance with MPSA measurements and lower specificity in recovering pathogenic variants. Some tools, including MMSplice and SQUIRLS, provide splicing effect predictions specific to all overlapping transcripts, and could permit investigation of isoform specific effects at the cost of reviewing many additional variant scores.

One limitation of our study is that the splicing assays we drew from made certain tradeoffs in exchange for scale. Minigene-based MPSAs necessarily include only minimal sequence context, and cannot capture effects from transcription elongation rate, or nucleosome positioning, each of which can influence splicing^97^. MPSA and SGE experiments typically use immortalized cancer cells, in which the splicing factor milieu may differ from that of the relevant tissues *in vivo*. Nevertheless, minigene assays are often well correlated across cell lines^2,6,14,48,53^ and have a sufficient track record of concordance with blood RNA analysis that they are often deemed acceptable as functional evidence during clinical variant interpretation^43,98,99^. Moreover, even when minigene assays misidentify a variant’s aberrant splicing outcome(s), they may still correctly flag the variant itself as splice disruptive^47,100^. In the future, improved splice effect benchmarking data could result from MPSAs with longer sequence contexts^101^ or delivered to more relevant tissues^102^ and from emerging approaches for in situ genome engineering^56,103^. Additionally, the growing usage of RNA-seq in genetic testing^23-25^ provides an opportunity to contribute both SDVs and neutral variants to the training and validation of future splice predictors.

## Conclusions

Here we have shown that saturation MPSAs provide an opportunity to critically evaluate the performance of computational splice effect predictors. Our results complement past benchmarking efforts using clinical variants and more broadly targeted MPSAs by testing algorithms’ ability to distinguish many variants’ effects within the context of several exons. This classification task resembles that faced by clinicians during variant interpretation, as there are many rare variants which do not impact splicing even in disease gene exons prone to splice disruption. We identified SpliceAI and Pangolin as the top-performing tools, but noted shortcomings including exonic variant performance, as well as practical challenges that end-users may encounter including selection of thresholds and the need for careful attention to gene model annotations. The continued growth of MPSA screens will present an opportunity to further improve splice effect predictors to aid interpretation of variants’ splicing impacts.

## Methods

### Saturation mutagenesis datasets

Splice effect measurements were obtained for a total of 3,616 variants in *POU1F1* (exon 2), *RON* (exon 11), *FAS* (exon 6), *WT1* (exon 9), *BRCA1* (11 exons) from the respective studies’ supplementary materials^48-50,53,56^. Variants were labeled as splice disruptive (SDV), intermediate, or neutral according to the classification made by each study; intermediate effect variants (*n*=121) were removed. The *RON* MPSA used a minigene spanning exons 10, 11, and 12, but as that assay did not measure skipping of exons 10 or 12, we only included variants most likely to influence exon 11 inclusion (i.e., within exon 11 and proximal halves of its flanking introns). Intronic variants more than 100 bp from either end of the selected exon were also discarded (*n*=93). *BRCA1* SGE measurements reflect both protein loss of function and mRNA effects, so we retained only synonymous and intronic variants so as to discard variants with effects that may be independent of splicing, and we further restricted to internal coding exons. For MPSAs in *POU1F1, RON*, and *WT1* that reported effects upon usage of multiple isoforms, we used for each variant the isoform score that was most highly significantly different than baseline (that is, maximum absolute z-score across isoforms per variant). For consistency in direction of effect (higher measured scores denoting greater splice disruption), *BRCA1* RNA and function scores’ signs were reversed. *FAS* enrichment scores were used without modification.

### Manual curation of clinical *MLH1* variants

A literature search was conducted for variants assayed for splicing effects in the tumor suppressor gene *MLH1*, yielding 77 publications (publication years 1995-2021). We included only single-base substitutions and required each variant’s splicing effects be supported either by RT-PCR and sequencing from patient blood-derived RNA or by mini-gene analysis. One exception is that essential splice site dinucleotide variants from Lynch Syndrome patients were included without molecular evidence, as loss of the native site would be considered strong evidence of pathogenicity by ACMG guidelines^26^. Any splicing outcome other than full exon inclusion was considered pathogenic^104^. Eight variants had conflicting reports (i.e., both pathogenic and benign) and were resolved with a majority vote among the reporting publications with ties being considered pathogenic. The final dataset included 296 variants (mean: 1.8 references per variant), of which 160 were splice disruptive.

### Random background variant set

We randomly drew 500,000 SNVs from within and near protein-coding genes to serve as a background set of exonic and proximal intronic variants with the potential to affect splicing. We used MANESelect canonical gene model annotations (version 1.0)^105^, restricting to protein coding transcripts with at least three coding exons. We discarded transcripts that had exons overlapping or within 100 bp of exon(s) of another transcript (on either strand), so that the variants’ classification (intronic vs exonic, proximity to splice site) would not depend upon the choice of gene model; this left 79.6% of all MANESelect transcripts (n=14,618/17,631). SNVs were selected at random from internal coding exons (padded by +/- 100 bp), and then these background SNVs were scored by splice effect predictors.

### Scoring with eight splice effect predictors

Pangolin version 1.0.2 was run with masking enabled and the distance parameter set equal to the length of the scored exon (for *MLH1* and *BRCA1*: the longest exon for each gene; for background set SNVs: 300 bp); reported Pangolin_max scores were used. SpliceAI version 1.3.1 was run via the python interface using a custom wrapper with masking enabled and distance setting following the same process as for Pangolin. For the transcriptomic background set, due to the high computational time to run SpliceAI, we downloaded version 1.3 precomputed scores from Illumina BaseSpace. SQUIRLS version 1.0.0 and MMSplice version 2.2.0 were both run on the command line with default settings to compute SQUIRLS score and delta logit PSI values respectively. HAL was run via the web interface (http://splicing.cs.washington.edu/SE) to predict exon skipping effects. HAL requires a baseline percent spliced in (PSI) value for the wildtype sequence (a parameter which has some predictive value on its own^38,95^). For this parameter, we used the following values: 90% for *MLH1*, 90% for *POU1F1*, 50% for *FAS*, 60% for *RON*, 80% for *BRCA1*, and 60% for *WT1*, based upon WT PSI values from the single exon MSPA original publications (rounded to the nearest 10%) and/or those exons known splicing patterns^104^. For background set exons, we used 50% to allow for HAL to predict the full range of outcomes from exon skipping (negative values) to increased inclusion (positive values), and to reflect the practical challenge in determining exon specific WT PSI values genome wide. Results with HAL were generally robust to the choices of WT PSI. For SPANR, S-Cap, and ConSpliceML, we obtained precomputed scores (SPIDEX zdelta PSI scores for SPANR; sens scores for S-Cap; ConSpliceML scores for ConSpliceML) from publicly accessible databases provided by the tools’ authors. For essential splice site dinucleotide scores, S-Cap provides two models (dominant and recessive), and we selected the lowest score (most severe predicted impact). We then transformed the S-Cap scores (taking 1-x, for input scores x in [0,1]) to match the direction of effect for other tools with higher values indicating greater likelihood of splice disruption.

We selected the MANESelect transcript model for each gene tested in the benchmarking set: ENST00000350375.7 (*POU1F1;* corresponding to the predominant isoform alpha), ENST00000452863.10 (*WT1* KTS+ isoform), ENST00000296474.8 (*RON*), ENST00000231790.8 (*MLH1*), ENST00000652046.1 (*FAS*), and ENST00000357654.9 (*BRCA1*). MMSplice, SQUIRLS, SpliceAI, and Pangolin all require an accompanying annotation file, and for a fair comparison, we provided each tool an identical annotation in which only the canonical transcript within the region of interest were included. Pre-computed ConSpliceML scores were selected by matching to the genomic position and relevant gene name. SQUIRLS’ annotation file was not readily customizable, so we used the default hg19 ENSEMBL annotation files that it supplies. We verified that at or near the tested exons, there were no differences between the selected gene models provided to other tools and the gene models within SQUIRLS’ annotations (ENST00000350375.2 for *POU1F1* alpha, ENST00000452863.3 for *WT1* KTS+, ENST00000296474.3 for *RON*, ENST00000231790.2 for *MLH1*, ENST00000355740.2 for *FAS*, ENST00000357654.3 for *BRCA1*). Substantive results did not change for the annotation dependent tools (MMSplice, Pangolin, SQUIRLS, and SpliceAI) when they were scored against annotation files including both alternative isoforms within *POU1F1* (beta isoform: ENST00000344265.8 for MMSplice, SpliceAI, and Pangolin and ENST00000344265.3 for SQUIRLS) and *WT1* (KTS-isoform: ENST00000332351.9 for MMSplice, SpliceAI, and Pangolin and ENST00000332351.3 for SQUIRLS). Within the testing using both alternative isoforms, for MMSplice and SQUIRLS, the most severe predicted impact from both isoforms was examined, and for Pangolin and SpliceAI, masking was based on both transcripts. For the transcriptomic background set, some variants either did not have a precomputed score for some tools, or the precomputed score entry mismatched the gene name or accession; this led to a small amount of missingness for some tools expected to score every SNV (SPANR: 1.6% of background variants excluded, SQUIRLS: 9.4%, SpliceAI: 4.1%). Pangolin and MMSplice each scored every background SDV. HAL only scores exonic variants, so all intronic variants were missing (56.5% of the background set), and S-Cap scores only some synonymous variants and variants within 50 bp of the splice sites, so 61.0% of background variants were missing.

### Variant classes

To examine performance within different gene regions, we categorized variants as those in: (i) essential splice site dinucleotides, (ii) intron near junction (3-10 bp from nearest exonic base), (iii) proximal intron (11-100 bp from nearest exonic base), (iv) exon near junction (<10 bp from nearest intronic base), and (v) deep exon (≥ 10 bp from nearest intronic base). For variants in multiple transcripts, the category with the most severe consequence was chosen (order: essential splice > exon near junction > intron near junction > deep exon > proximal intron). We assessed the abundance of each variant class within previously curated clinical variant sets. For the S-Cap training set we combined the proportions of 5’ core, 5’ core extended, and 3’ core variants listed from their clinically derived pathogenic set in Figure 1C^29^, and for SQUIRLS we tallied variant classes across training data without alterations^30^.

### Nominating annotation-sensitive alternatively spliced genes

To identify genes with alternative splice forms for which choice of annotation could influence splicing effect predictions, we obtained exon-exon junction read counts from GTEx portal (version 8). We restricted to protein coding genes (*n*=19,817) and computed, for each of 54 tissues, the median junction read counts per million junction reads (junction CPM) across samples of that tissue for junctions that fell within coding portions of their respective genes. Junctions with a junction CPM ≥ 0.1 were considered expressed (*n*=14,831 genes had at least one expressed junction in at least one tissue). Next, we identified 12,124 genes where at least one expressed splice site was alternatively used in multiple junctions. Within each group of alternative junctions at a given splice site (e.g., two junctions corresponding to one donor paired with either of two different acceptors corresponding to skipping or inclusion of a cassette exon), we computed the fractional proportions of each junction’s use and determined which alternate junctions were included in SpliceAI’s default annotations. Fractional proportions were computed separately for each tissue. We deemed ‘moderately used unannotated’ splice sites as any group of alternative junctions with at least one unannotated expressed junction which had ≥20% fractional usage in a given tissue. As a specific example of an exon sensitive to annotation selections, we scored *FGFR2* exon 8 (also called exon IIIc) against two alternative isoforms (ENST00000358487.10 for FGFR2c and ENST00000358487.5 for FGFR2b (default)) using our custom SpliceAI wrapper with masking turned on and a scoring distance equal to the length of the exon.

### Statistical methods

Area under the curve metrics were calculated using sklearn in python. For **Figure 2**, a single cutoff was selected for each tool that maximized Youden’s J for identifying SDVs across the full *POU1F1* and *WT1* MPSA variant sets, respectively. Tools with fewer than ten scored variants in each pre-defined region were excluded. To compute transcriptome-adjusted sensitivity for each algorithm, we first computed the score threshold *t*(*x*) at which that algorithm called a given fraction *x* of the transcriptomic background set as disruptive (for all values of *x* in [0,1]). Transcriptome-adjusted sensitivity was then the sensitivity for benchmark SDV detection at this threshold: (# benchmark SDVs with score≥t(x)) / (# benchmark SDVs). Area under the curve was then taken for transcriptome-adjusted sensitivity as a function of the background set fraction *x*, and was computed using the sklearn auc function.

To analyze correlation between algorithms and MPSA measurements, the absolute value of each algorithm’s score was taken to accommodate HAL, MMSplice, SPANR, and Pangolin for which signed scores indicates exon skipping vs inclusion. *FAS* was one exception to the rule: since *FAS* enrichment scores directly measured exon skipping (negative values) and exon inclusion (positive values), for signed scoring tools (HAL, MMSplice, SPANR, and Pangolin) we compared signed *FAS* scores with tools’ signed values, and for the rest (SpliceAI, SQUIRLS, ConSpliceML, S-CAP), compared absolute values of the measured scores with the tools’ scores. For classification performance analyses (prAUC, transcriptome-adjusted sensitivity), absolute values of tools’ predicted scores were used.

## Acknowledgements

Datasets scored by all the algorithms are available as Supplementary Tables, and the Jupyter notebooks needed to reproduce the scoring and plots are posted at GitHub.

## Acknowledgements

We thank Sally Camper, Jen Lai Yee, and members of the Kitzman lab for helpful discussions.

## Author contributions

C.S. analyzed the data. C.S. and J.O.K. wrote the manuscript.

