## Supplementary Figures for "Benchmarking splice variant prediction algorithms using massively parallel splicing assays"

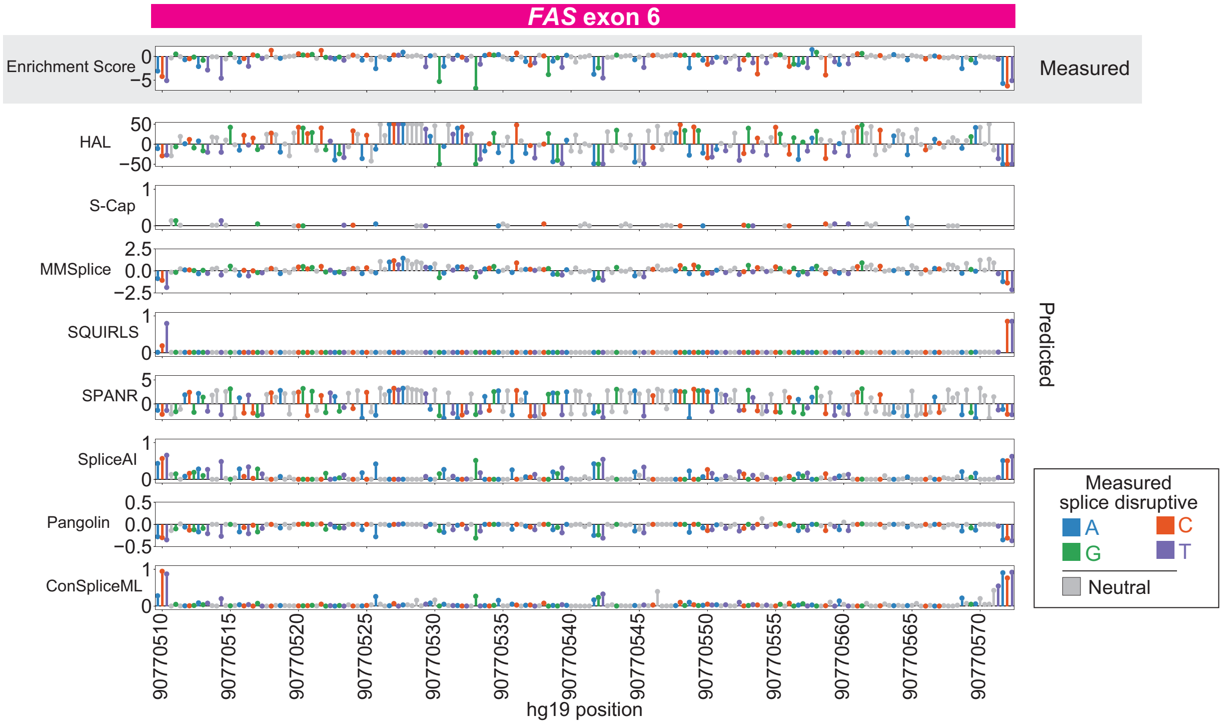


**Supplementary Figure 1. Splicing effect map and bioinformatic predictions for *FAS* exon 6.** MPSA measured enrichment score of *FAS* exon 6 (gray, top panel; increased skipping – negative values, increased inclusion – positive values), along with bioinformatic predictions (subsequent panels), with splice effects/predictions plotted by variant position. Each lollipop denotes one variant, shaded by effect in MPSA (gray: neutral, colors: SDVs, shaded by mutant base).

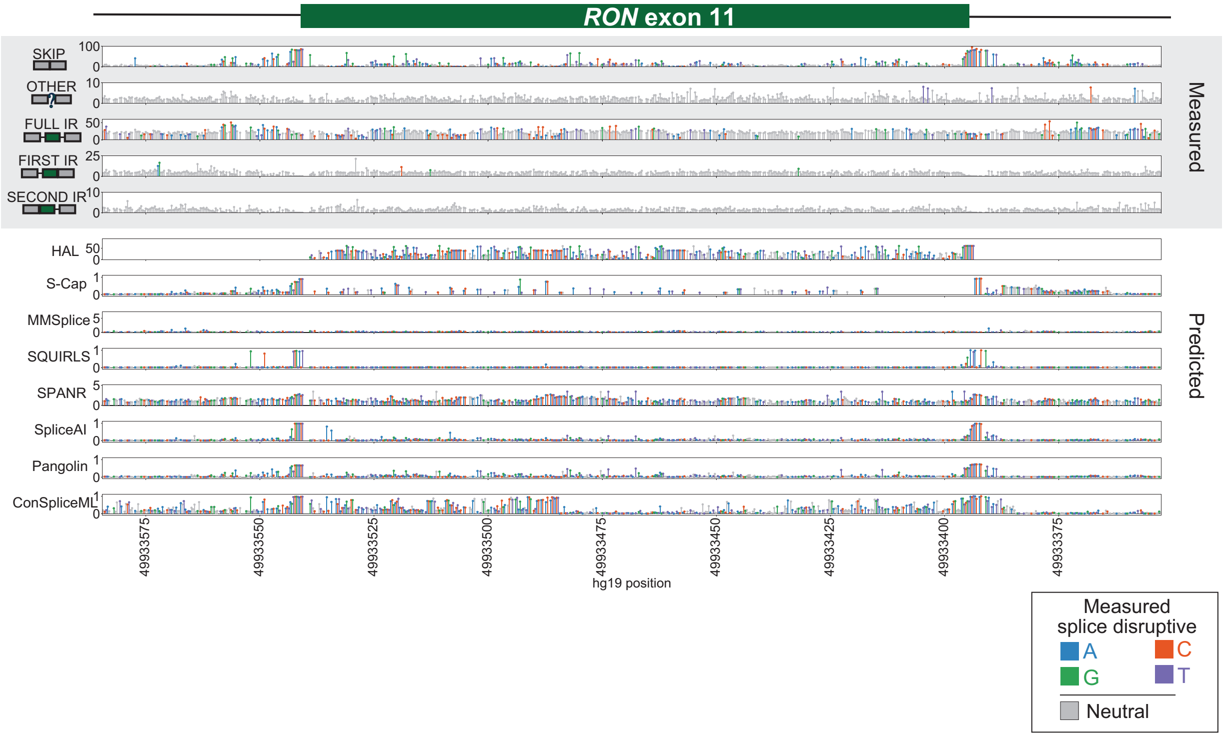


**Supplementary Figure 2. Splicing effect map and bioinformatic predictions for RON exon 11.** MPSA measured percent usage for different splicing outcomes at RON exon 11 (gray, top panel): skipping, other isoforms, full intron retention ("FULL IR"), first intron retention ("FIRST IR"), and second intron retention ("SECOND IR"), along with bioinformatic predictions (subsequent panels), with splice effects/predictions plotted by variant position. Each lollipop denotes one variant, shaded by effect in MPSA (gray: neutral, colors: SDVs, shaded by mutant base).

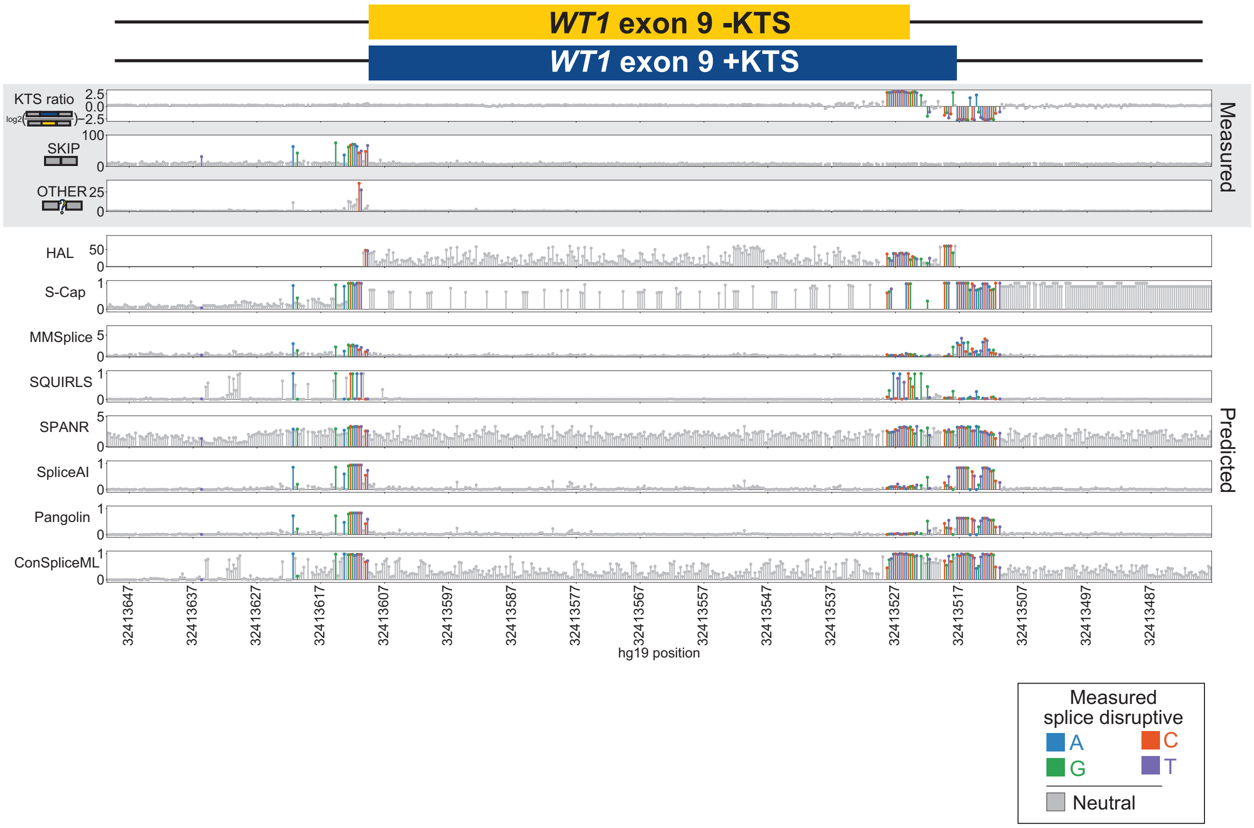


**Supplementary Figure 3. Splicing effect map and bioinformatic predictions for *WT1* exon 9.** MPSA measurements for different splicing outcomes at *WT1* exon 9 (gray, top panel): log2ratio(%KTS+/%KTS-), percent exon skipping, and percent other isoforms, along with bioinformatic predictions (subsequent panels), with splice effects/predictions plotted by variant position. Each lollipop denotes one variant, shaded by effect in MPSA (gray: neutral, colors: SDVs, shaded by mutant base).

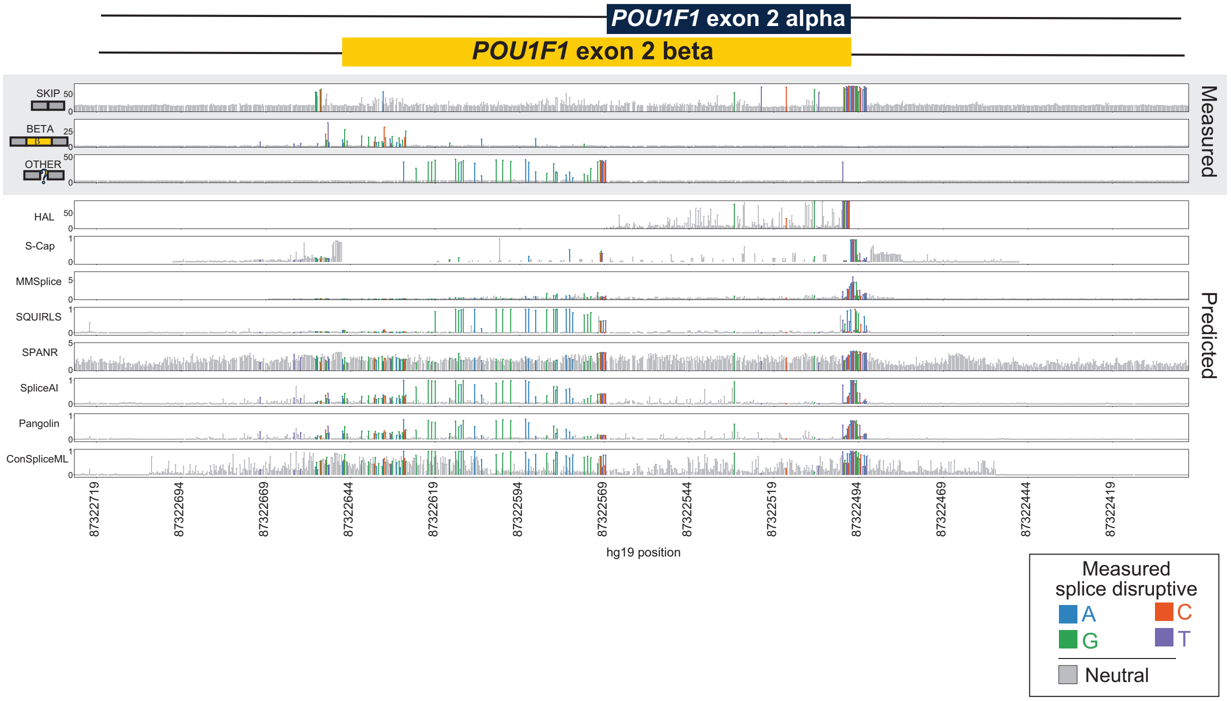


**Supplementary Figure 4. Splicing effect map and bioinformatic predictions for POU1F1 exon 2.** MPSA measured percent usage for different splicing outcomes at POU1F1 exon 2 (gray, top panel): exon skipping, exon 2 beta, and other isoforms, along with bioinformatic predictions (subsequent panels), with splice effects/predictions plotted by variant position. Each lollipop denotes one variant, shaded by effect in MPSA (gray: neutral, colors: SDVs, shaded by mutant base).

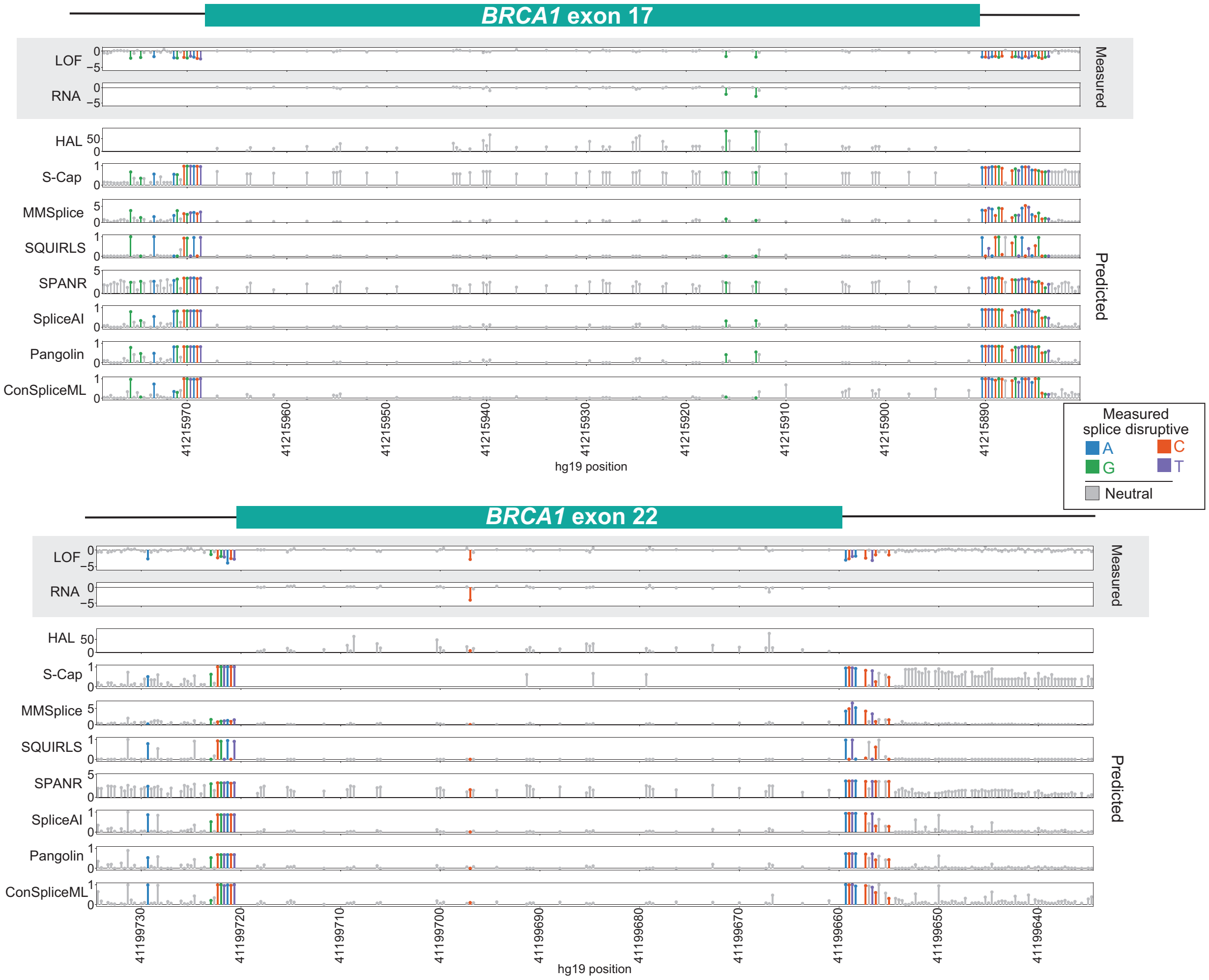


**Supplementary Figure 5. Splicing effect map and bioinformatic predictions for *BRCA1* exons.** SGE measurements for two representative *BRCA1* exons (exons 17 and 22; in gray, top panel): log2-ratio function score and log2ratio RNA score, along with bioinformatic predictions (subsequent panels), with splice effects/predictions plotted by variant position. Each lollipop denotes one variant, shaded by effect in SGE (gray: neutral, colors: SDVs, shaded by mutant base). Missense and stop gained variants are excluded.

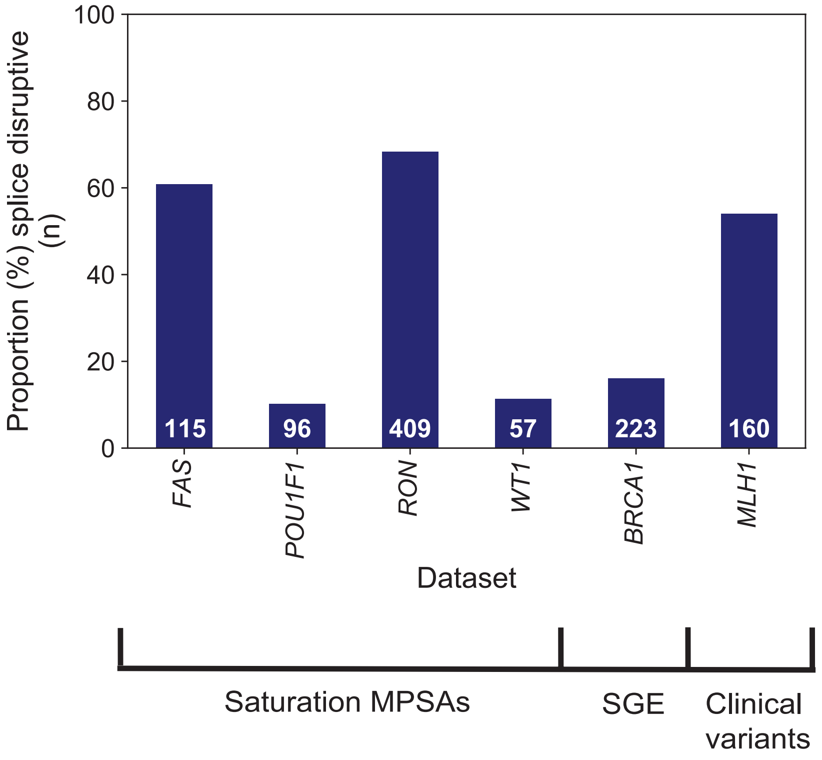


**Supplementary Figure 6. Proportion of splice disruptive variants (SDVs) within benchmarked datasets.** Bar plot showing the proportion of SDVs out of all measured variants (y-axis) within the saturation MPSAs (*FAS*, *POU1F1*, *RON*, *WT1*), SGE (*BRCA1*), and clinically curated variant set (*MLH1*) (x-axis). Numbers on each bar display the count of splice disruptive SNVs per dataset.

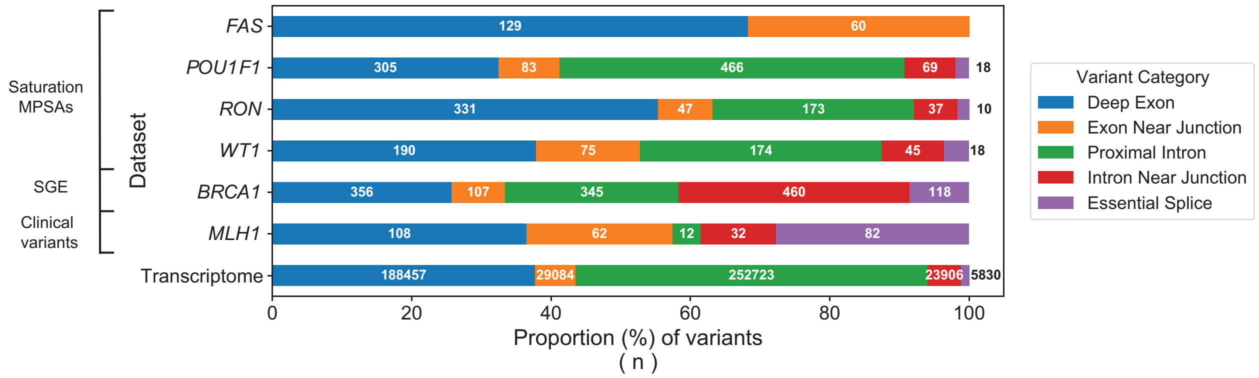


**Supplementary Figure 7. Breakdown of benchmark and background variant sets by variant class**. Proportions (x-axis) of variant category (color) deep exon (blue) in each benchmark variant dataset. Datasets are grouped by study type (MPSAs, SGE, and clinical variants), and ‘transcriptome’ denotes the random background set of variants.  Variant categories are defined and shaded as in **Figure 1B**. Numbers of each bar indicate count of each type of variant per dataset.

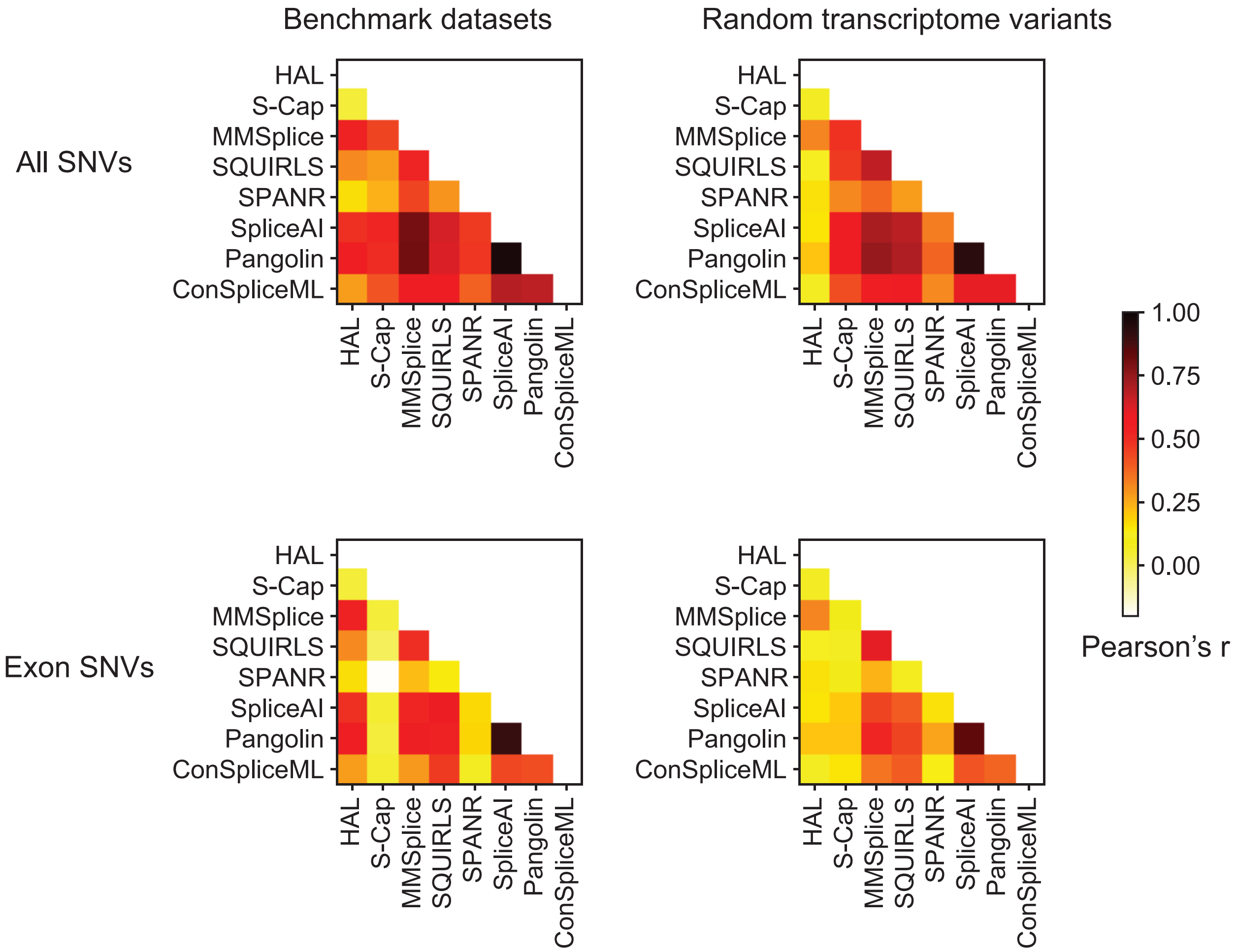


**Supplementary Figure 8. Correlations among bioinformatic algorithms.** Heatmaps of Pearson correlations between scores from eight bioinformatic algorithms across benchmarked variants (left column) and randomly selected 'background set' variants (right column). Top row shows correlations across all variants; bottom row shows correlations over only exonic variants.

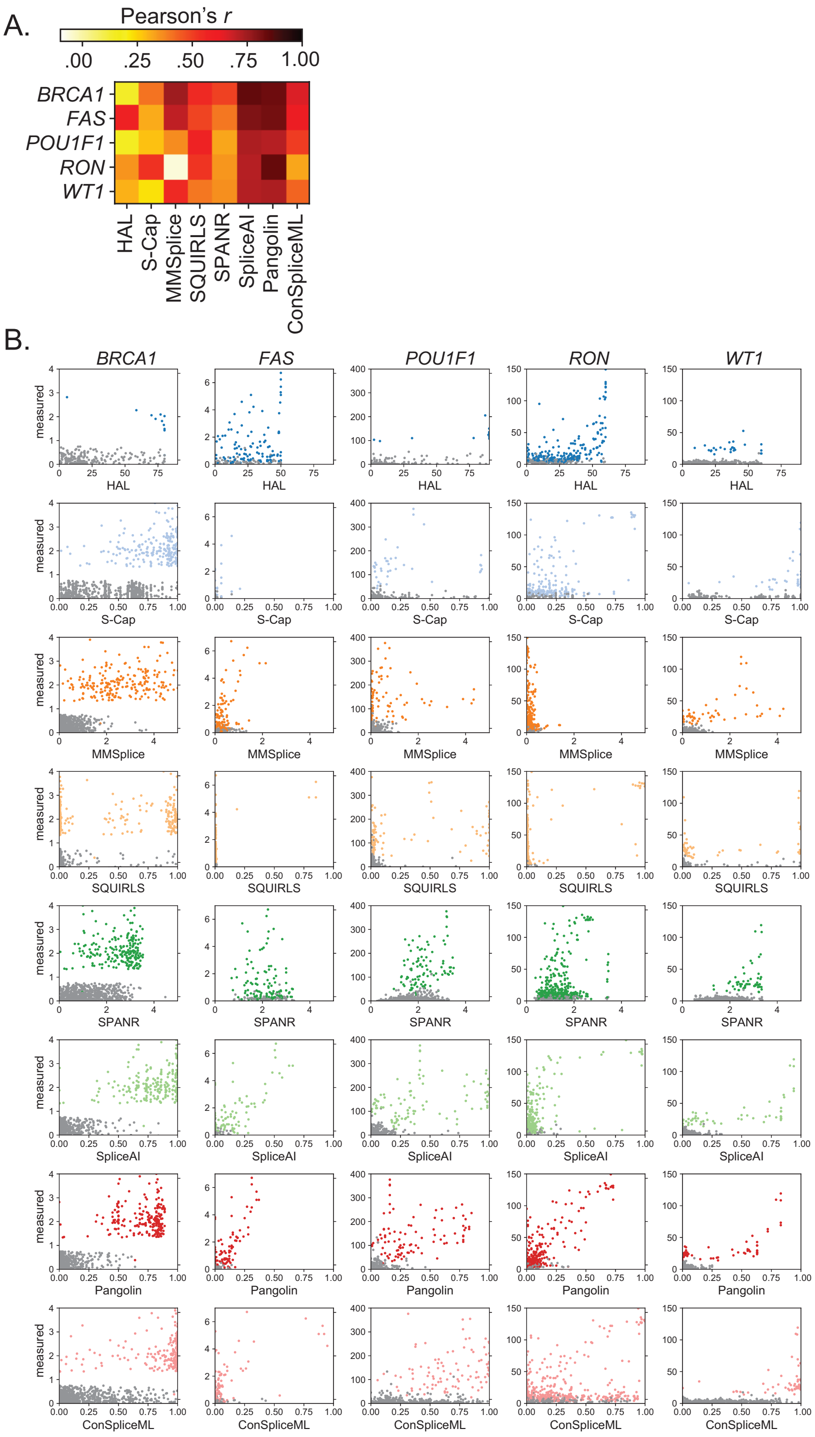


**Supplementary Figure 9. Correlations between bioinformatic algorithms' scores and MPSA measurements. (A)** Heatmap showing the Pearson's correlations between bioinformatic algorithms (x-axis) and MPSA-measured effects (y-axis). *MLH1* SNVs are omitted as they were curated across many different studies and do not have measurements beyond classification as deleterious/neutral. **(B)** Scatterplot of all variants showing measured effect (y-axis) and predicted effect (x-axis); gray points denote splice neutral variants and shaded points are splice-disruptive variants.

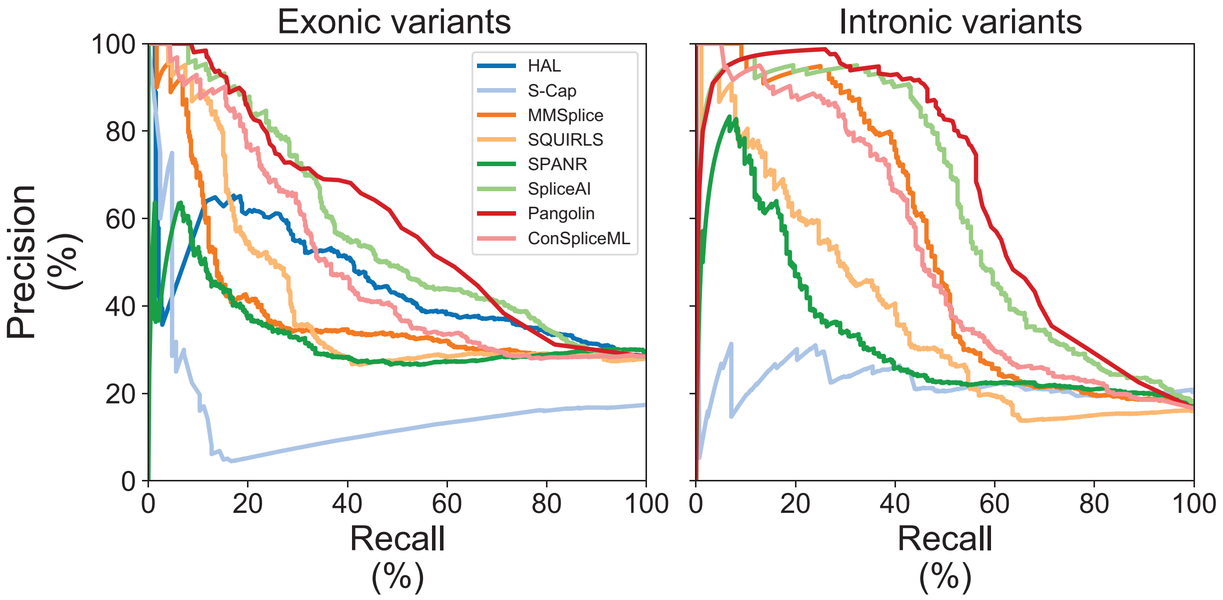


**Supplementary Figure 10. Classification performance without essential splice site mutations.** Precision-recall curves showing algorithms' performance at distinguishing SDVs from splicing-neutral variants in each dataset, for exonic variants (identical to **Figure 3C**) and intronic variants after removing variants at essential splice sites.

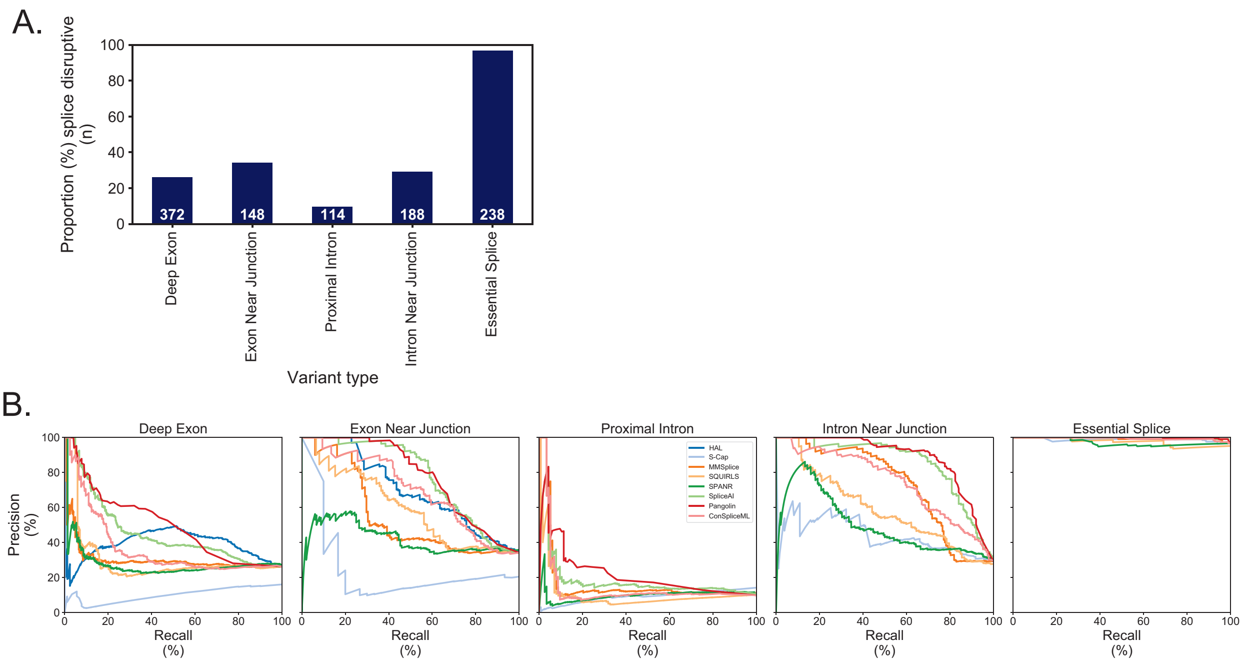


**Supplementary Figure 11. Classification performance by variant category. (A)** Proportion of variants which are splice disruptive by variant category. Counts of splice disruptive variants in each category are inset. **(B).** Precision-recall curves showing algorithms' performance at distinguishing SDVs from splicing-neutral variants in each variant category as defined in **Figure 1B**.

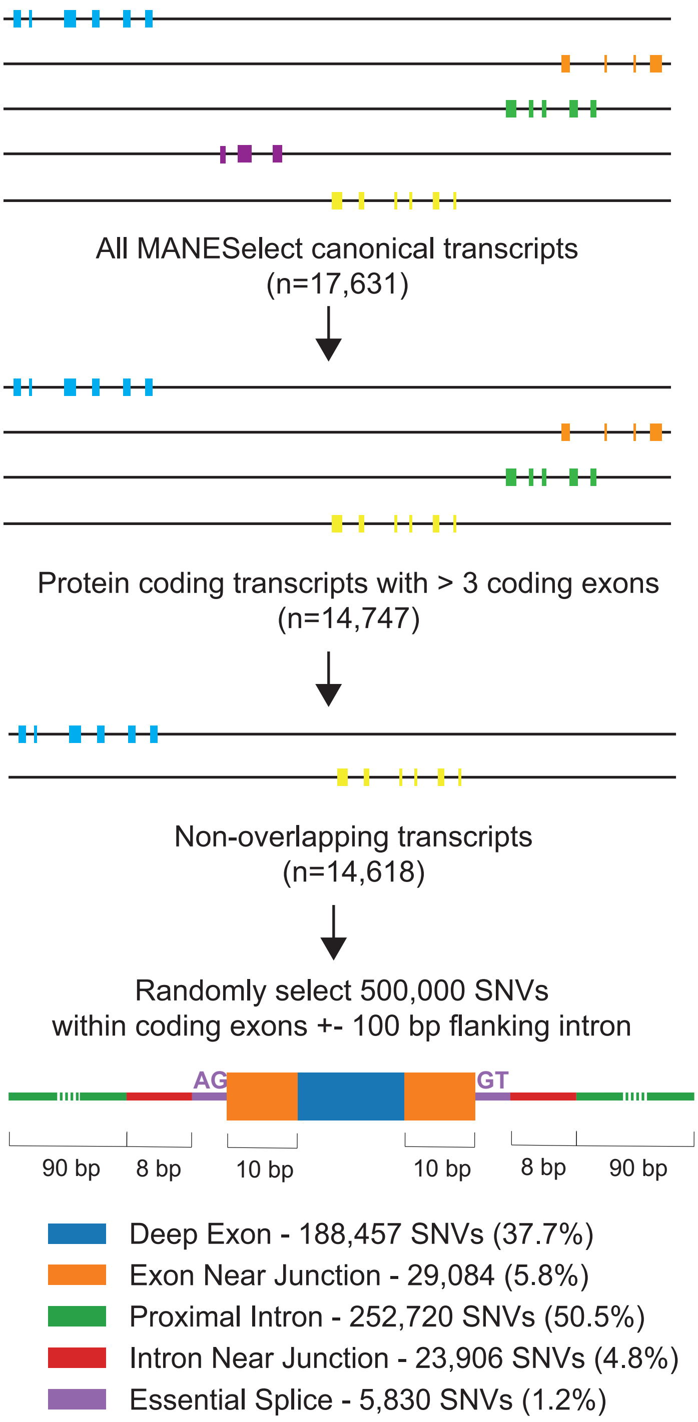


**Supplementary Figure 12. Background set of random exonic and near-exonic variants.** Schematic shows criteria used to select gene models and counts of MANESelect transcripts remaining at each step. At bottom, counts and proportions of background set variants by category.

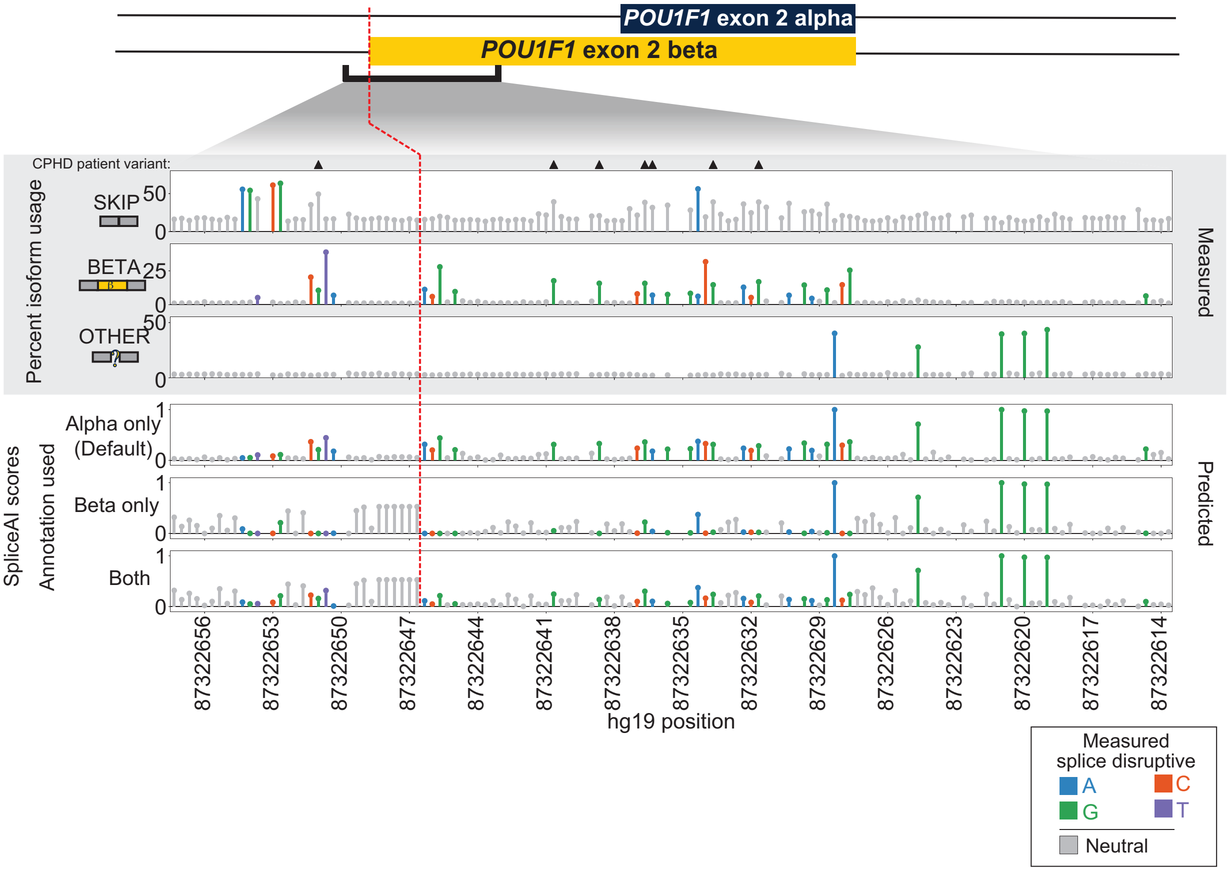


**Supplementary Figure 13. Effects of gene model annotation on SpliceAI predictions near *POU1F1* exon 2 beta acceptor.** MPSA measured percent usage of *POU1F1* isoforms are shown in the upper tracks (gray background), with variants called SDVs shaded with color and denoted as in **Supplementary Figure 4**. SpliceAI deltaMax scores are shown in the bottom three tracks, obtained using default annotation (alpha isoform only; top), beta isoform only (middle), or both isoforms (bottom). Combined pituitary hormone deficiency (CPHD) patient variants are marked with black triangles.

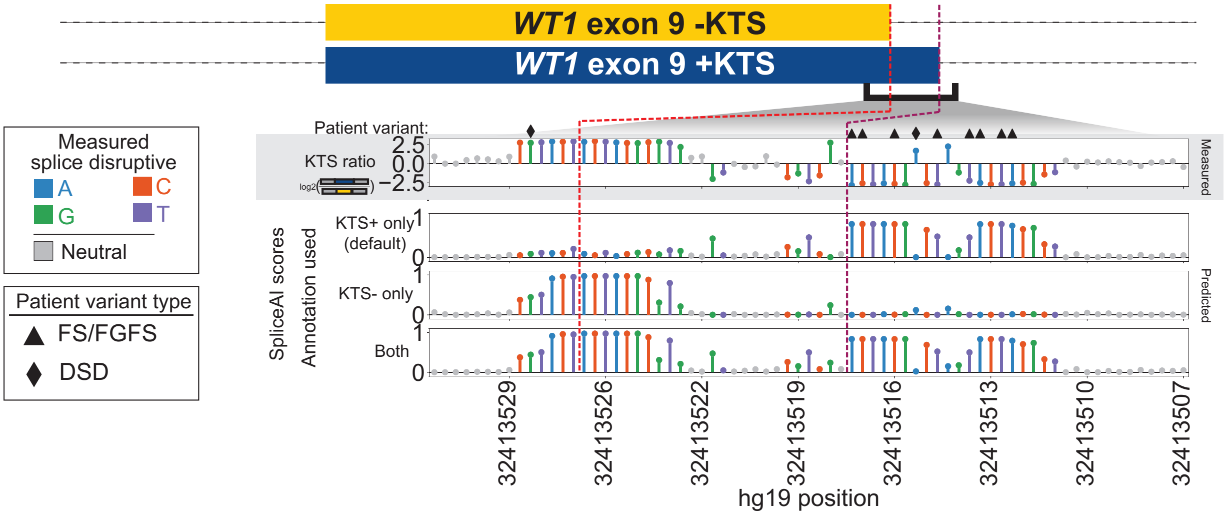


**Supplementary Figure 14. Effects of gene model annotation on SpliceAI predictions at *WT1* exon 9 alternate donors.** MPSA measured $\log_2$ratio of *WT1* exon 9 alternate isoforms (top panel) along with masked SpliceAI predictions scored with annotation files of the KTS+ isoform (second panel), KTS- isoform (third panel), and both isoforms (bottom) by variant position (x-axis). Gray lollipops denote MPSA measured splicing-neutral variants, while shaded lollipops indicate the base pair change of each measured SDV (dark colors). Pathogenic Frasier's syndrome (FS) and focal segmentation glomerulosclerosis (FSGS) clinical variants are shown with black triangles, and variants observed in individuals with 46,XX ovotesticular differences in sexual development (OTDSD) are denoted by black diamonds.

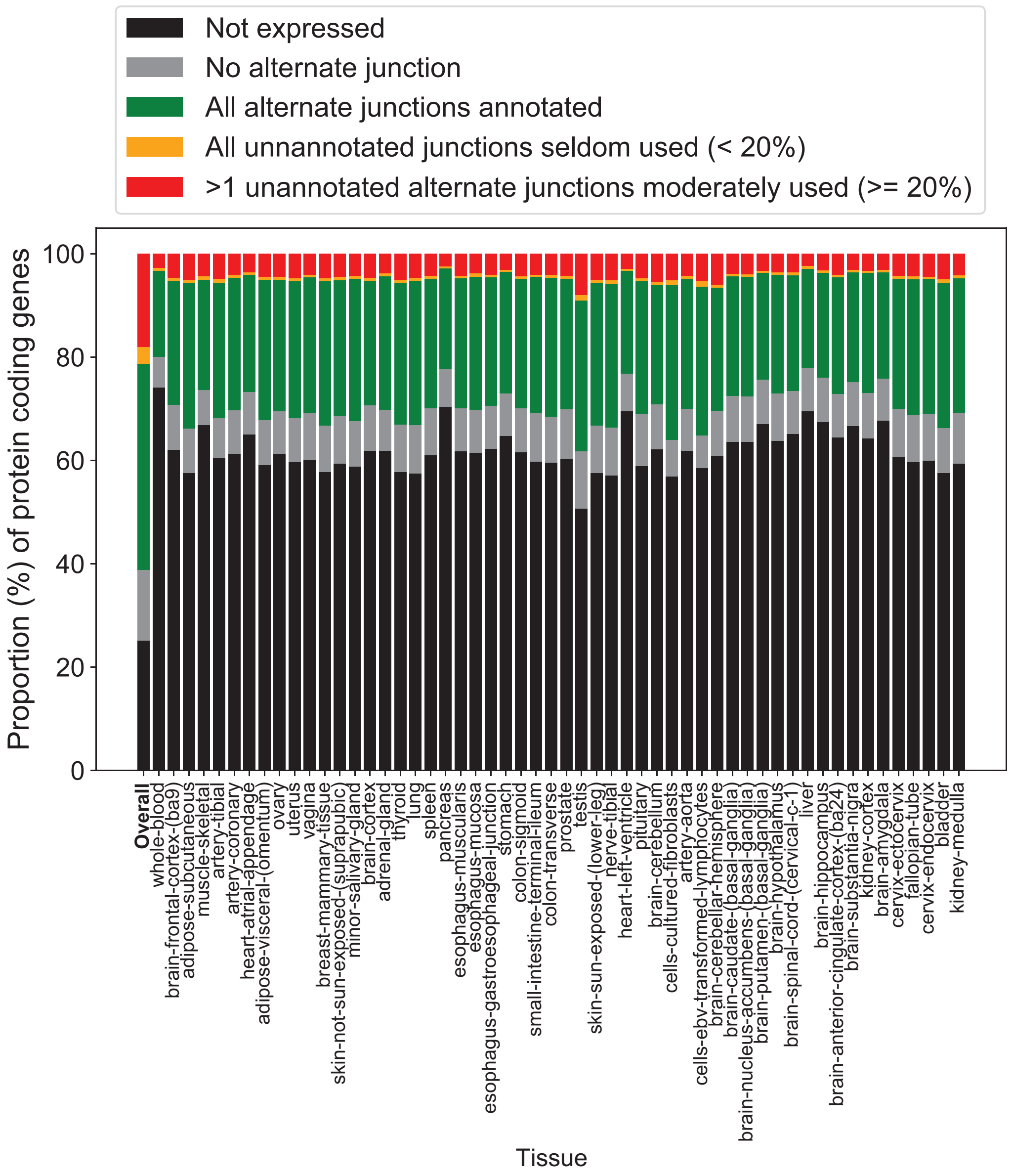


**Supplementary Figure 15. Annotation sensitive alternatively spliced genes.** Proportion of protein coding genes within GTEx (y-axis) that are either not expressed (CPM < 0.1; black), have no expressed alternate splice junctions (gray), have all alternatively used splice junctions present in SpliceAI annotations (green), have only seldom used unannotated alternate splice junctions (orange), or have at least one unannotated alternate splice junction with at least modest use (≥ 20%; red). Proportions are shown across all tissues (first bar labeled ‘overall’) and within individual tissues.

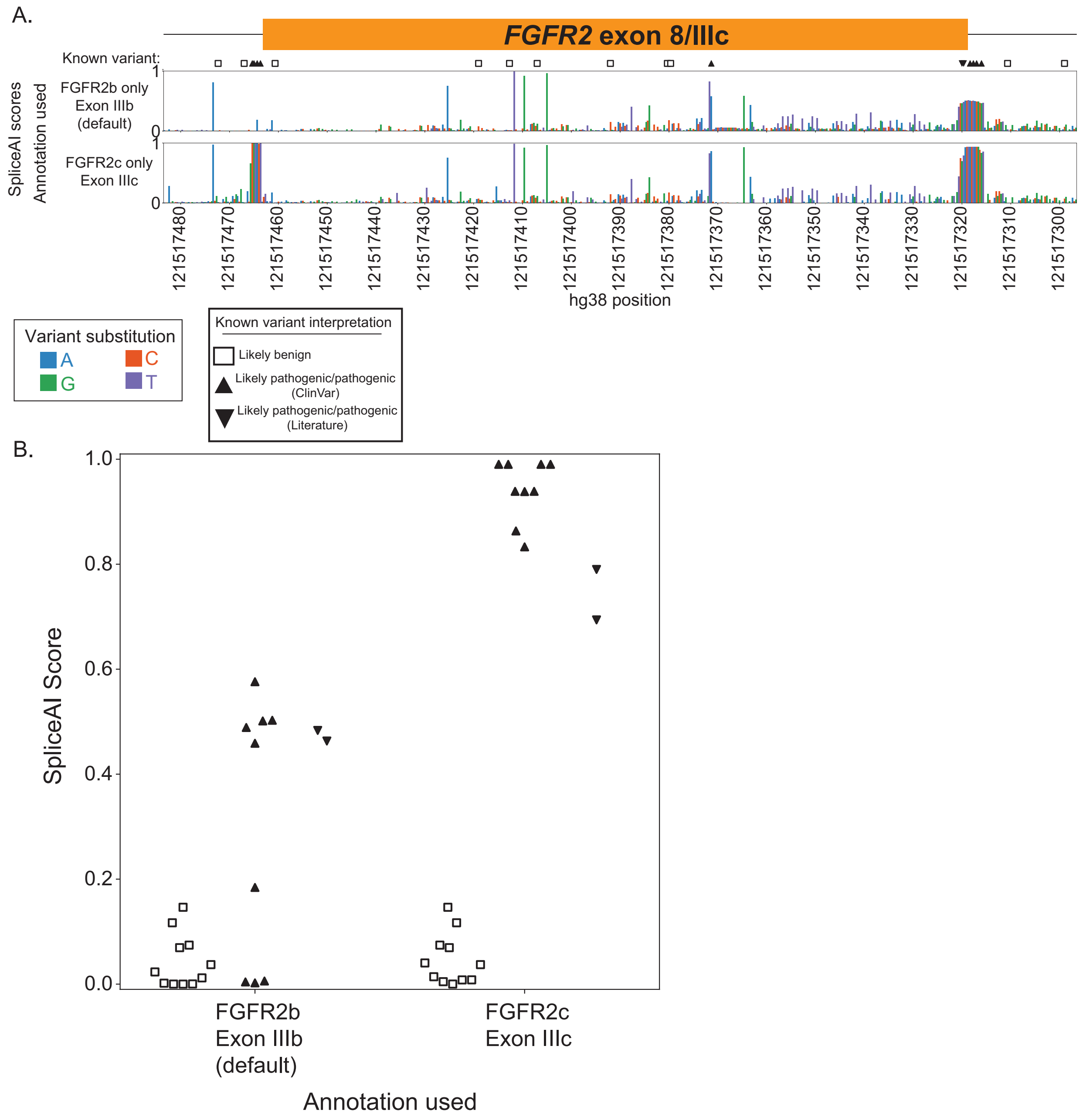


**Supplementary Figure 16. Effects of annotation changes within SpliceAI for FGFR2 exon IIIc. (A)** Tracks of SpliceAI scores (y-axis) for all SNVs in *FGFR2* exon IIIc showing either score using masking and default annotation (upper panel) or scores with masking and only the FGFR2c isoform (lower panel) vs hg38 position (x-axis). Bars are colored by their nucleotide substitution. Symbols denote synonymous and intronic likely benign (empty square) or likely pathogenic/pathogenic variants (black triangle) from ClinVar or published reports (inverted black triangle). Several pathogenic exon IIIc acceptor disrupting variants are missed when annotation does not include exon IIIc **(B)** SpliceAI scores (y-axis) vs annotation used (x-axis), each marker indicates published or ClinVar interpretation as in A. Scores of pathogenic and benign variants in exon IIIc are perfectly separated when exon IIIc is included but not using default annotations.

**Supplementary Table 1.** Measured splicing scores and splicing outcomes as well as predicted scores for each of the eight algorithms evaluated for the six benchmarked datasets. Predicted scores from the eight bioinformatic tools for each of the 500,000 randomly selected exonic and near-exonic background variants. SpliceAI predictions against the FGFR2b and FGFR2c annotations for *FGFR2* exon IIIc.

**Supplementary Table 2.** Tool specific thresholds at which 5%, 10%, and 20% of the 500,000 background variants are predicted to be splice disruptive. Transcriptomic normalized sensitivity values for each algorithm and benchmarked dataset overall, within exonic variants, within intronic variants, and within intronic variants after removing variants at essential splice sites. Tables of sensitivity values are provided at transcriptome normalized cut offs of 5%, 10%, and 20% as well as the area under the curve (AUC) values.

**Supplementary Table 3.** Optimal score thresholds for each tool within the benchmarked datasets and across variant classes as defined in **Figure 1B**.
